# The role of olivary phase-locking oscillations in cerebellar sensorimotor adaptation

**DOI:** 10.1101/2024.03.06.583676

**Authors:** Niceto R. Luque, Richard R. Carrillo, Francisco Naveros, Eduardo Ros, Angelo Arleo

## Abstract

The function of the olivary nucleus is key to cerebellar adaptation as it modulates long term synaptic plasticity between parallel fibres and Purkinje cells. Here, we posit that the neural dynamics of the inferior olive (IO) network, and in particular the phase of subthreshold oscillations with respect to afferent excitatory inputs, plays a role in cerebellar sensorimotor adaptation. To test this hypothesis, we first modelled a network of 200 multi-compartment Hodgkin-Huxley IO cells, electrically coupled via anisotropic gap junctions. The model IO neural dynamics captured the properties of real olivary activity in terms of subthreshold oscillations and spike burst responses to dendritic input currents. Then, we integrated the IO network into a large-scale olivo-cerebellar model to study vestibular ocular reflex (VOR) adaptation. VOR produces eye movements contralateral to head motion to stabilise the image on the retina. Hence, studying cerebellar-dependent VOR adaptation provided insights into the functional interplay between olivary subthreshold oscillations and responses to retinal slips (i.e., image movements triggering optokinetic adaptation). Our results showed that the phase-locking of IO subthreshold oscillations to retina slip signals is a necessary condition for cerebellar VOR learning. We also found that phase-locking makes the transmission of IO spike bursts to Purkinje cells more informative with respect to the variable amplitude of retina slip errors. Finally, our results showed that the joint action of IO phase-locking and cerebellar nuclei GABAergic modulation of IO cells’ electrical coupling is crucial to increase the state variability of the IO network, which significantly improves cerebellar adaptation.

**Author summary:** This study aims to elucidate the dual functionality of the inferior olive (IO) in cerebellar motor control, reconciling hypotheses regarding its role as either a timing or instructive signal. Specifically, we explore the role of subthreshold oscillations (STOs) within the IO, investigating their potential influence on the climbing fibres-to-Purkinje cell spike pattern responses and subsequent cerebellar adaptation, notably during the vestibulo ocular reflex. Aiming these objectives, we constructed a detailed olivary network model within a cerebellar neural network, enabling a mechanistic analysis of the functional relevance of STOs in spike burst generation, propagation, and modulation within target Purkinje cells. Our findings reveal the intricate nature of complex spike bursts triggered by climbing fibres—IO axons—into Purkinje cell dendrites, demonstrating a hybrid nature involving binary clock-like signals and graded spikelet components acting as an instructive signal.

## I. Introduction

The role of the inferior olive (IO) in cerebellum-dependent motor control remains partially understood. IO is supposed to instruct a signal that triggers the associative synaptic plasticity at parallel fibres - Purkinje cell (PC) synapses [1]. IO is also hypothesised to provide a timing signal that drives downstream PC outputs thanks to its membrane potential subthreshold oscillations (STOs) [2-4]. Yet, there is a need to understand the role of STOs in the generation of IO spiking patterns transmitted to PCs through climbing fibres, (CFs). It is also necessary to study how STOs determine the neural state of the IO network and, ultimately, cerebellar adaptive dynamics.

Neighbour IO neurons dendritically contact each other in a glomeruli through which they are electrically coupled via gap junctions [5]. This electrical coupling enables the propagation of neural activity across the olivary network. IO internal conductance dynamics generate STOs [6], whose phase is assumed to determine spike burst responses to excitatory inputs [7, 8]. The STO phase-dependent gating mechanism can grade the spike burst lengths according to IO input amplitude, thus allowing more than binary “all-or-nothing” patterns to be encoded [9]. Furthermore, inhibitory inputs to the IO glomeruli from the medial cerebellar nuclei add another piece on the spike bursts generation and propagation jigsaw. These GABAergic synapses are known to modulate IO gap junctions, by reducing the electrical coupling and thus the synchrony amongst IO cells [10]. Therefore, IO drives PC complex spikes by weighting its inhibitory and excitatory inputs, which determines the IO neural activation and synchrony via STO phase-dependent modulation. This complex mechanism raises the question of how information is processed and transmitted by the IO network to facilitate cerebellar adaptation and motor control.

To address this question, we simulated a realistic IO network, and we incorporated it within a spiking cerebellar network. We then used the resulting feedforward control loop system to learn a specific sensorimotor adaptation task: the prediction of oculomotor commands for the acquisition of the vestibulo-ocular reflex (VOR). VOR counter rotates the eyes with respect to head rotations to stabilise the images on the retina, thus maintaining the image in the centre of the visual field. VOR has been profusely used as a model system to test the possible cerebellar role in motor learning [11] and feed-forward control adaptation [12]. The model presented in this study aims at investigating the functional relevance of olivary STOs in terms of spike burst generation, propagation and modulation of PC complex spikes, electrical coupling role on IO neural coding, as well as the interplay of all those mechanisms during VOR adaptation.

In particular, we postulate that within the olivary system, STOs may serve as neural pattern encoders. STOs, acting as a master clock within the IO, are phase-locked to the retinal slip signals, thereby finely regulating the neural response timing for cerebellar motor adaptation. We also test the hypothesis of a dual role of the IO, serving as both a master clock through STOs and as a graded instructive signal during VOR adaptation. Importantly, the complex spike (CS) bursts triggered by CF into PC dendrites exhibit a hybrid nature, combining binary and graded spikelet components. Additionally, we investigate the intricate interplay amongst inhibitory (GABAergic) and excitatory inputs, as well as electrical coupling within the IO network, shaping IO neural coding. The modulation of graded CS bursts depending on the retinal slip amplitude does not require GABAergic action to decrease IO electrical coupling and thereby disrupt olivary network synchronicity [13, 14]. Yet, we study whether the GABAergic desynchronising action of the olivary network may play a role in improving rotatory-VOR (r-VOR) adaptation.

## II. Results

### A. Olivary neural dynamics

#### 1. IO spike burst responses to excitatory dendritic inputs

We implemented each IO cell as a Hodgkin-Huxley model with 3 compartments: somatic, axonal, and dendritic (Fig 1A left; see Methods). The neuronal IO model reproduced the spike burst activity of real olivary cells in response to dendritic step current injections (Fig 1A centre) [15, 16]. It also captured the linear relation between the number of burst spikes and the amplitude of the excitatory synaptic input current (Fig 1A right).

**Fig 1.**
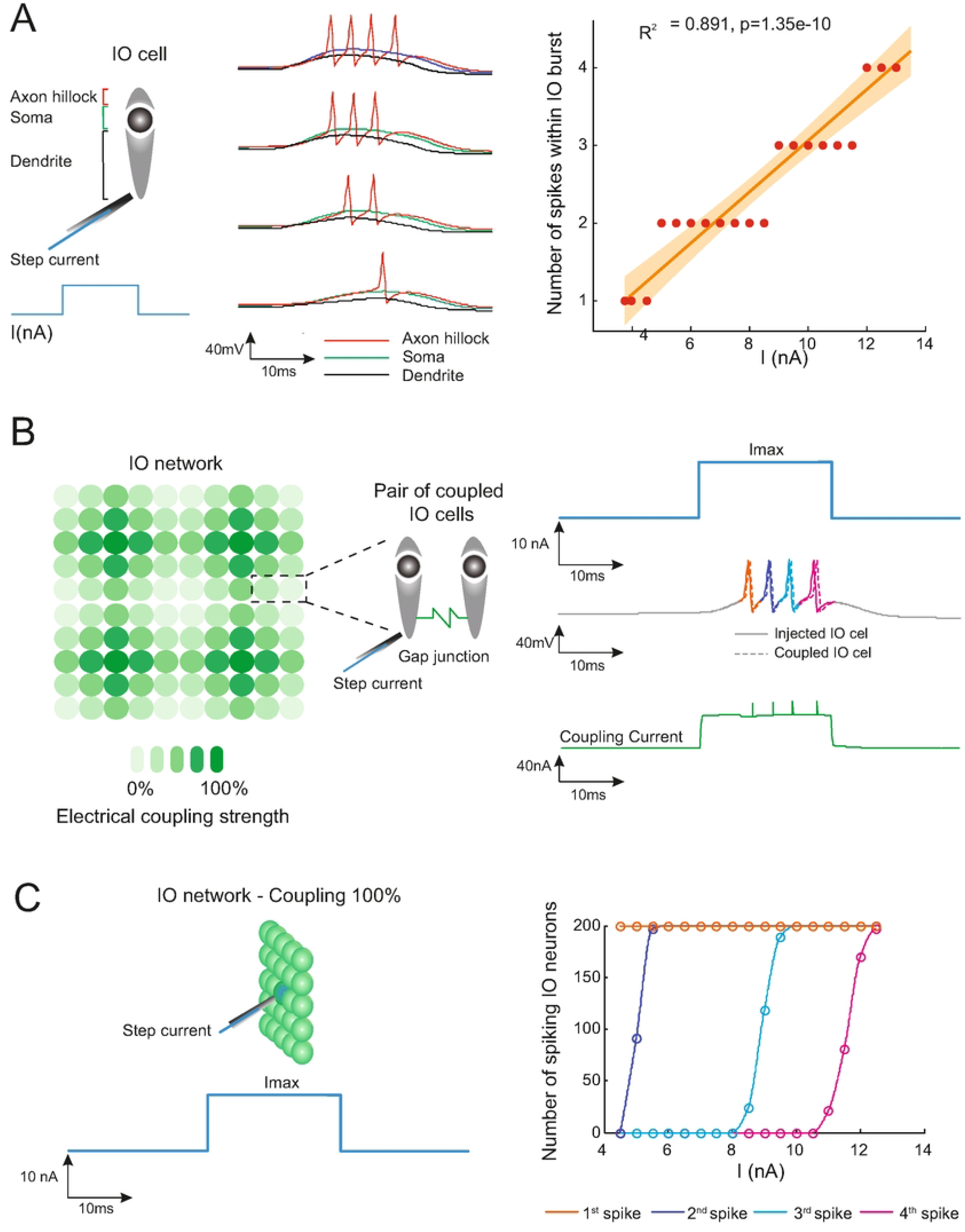
Electrophysiological properties of the HH IO model and the olivary network. **(A)** Schematic representation of the three-compartmental HH IO model used. The compartments represent the axon hillock, the soma, and the dendrite. To modulate the spike burst, a depolarising step current is applied to the dendritic compartment (black line), which exhibits a slow depolarization. The somatic compartment (green line) responds with a slow depolarization, whilst the axon hillock (red line) exhibits fast sodium responses to the somatic depolarisation, resulting in a burst of spikes. Regulating the amplitude of the depolarising step current applied to the dendritic compartment allows for the modulation of the number of spikes within the burst experienced by the axon hillock. The number of spikes within the burst increases linearly with the amplitude of the depolarising step current applied (R2: 0.891, p < 1 x 10^^^−10), enabling a graded codification that goes beyond the all-or-nothing IO learning paradigm [9, 18]. **(B)** The left-hand side plot depicts an IO network consisting of 200 three-compartment Hodgkin-Huxley (HH) inferior olive (IO) neurons arranged in a 3D lattice configuration with dimensions of 10 x 10 x 2 microzones. Each olivary microzone is a 10 x 10 x 1 subunit, and they are electrically coupled via gap junctions. The figure illustrates the electrical connectivity scheme for the 100 IO neurons within each microzone. The colour grading in the figure represents gap junctional interconnections within the lattice arrangement, where each neuron in the model is electrically connected only to its closest neighbours. The right-hand side plot depicts an example of axon hillock membrane potential traces during burst propagation due to IO-to-IO electrical coupling. A 10nA depolarising step current is applied for 20ms to the dendritic compartment of the left-hand side IO whilst maintaining maximal coupling strength, i.e., fully open gap junctions. The transmitted dendritic depolarised current via electrical coupling ensures the entire burst propagation (axon hillock membrane potential) to the right-hand side IO neuron in almost no time (microsecond scale). **(C)** IO neurons located at the centre of each 5 × 5 square within the lattice arrangement receive an instructive input signal. This input signal simultaneously reaches a subset of 5×5 neurons each time we simulate the activation of glutamate receptor channels. Burst propagation within the entire network depends on the effectiveness of the coupling. A graded [0-14nA] depolarising step current, applied for 20ms, reaches the dendritic compartment of the central IO neuron in all subsets of 5×5 neurons in the lattice olivary network, maintaining 100% effective coupling. Varying the current amplitude modulates the length of the burst. A 14nA depolarising step current, when coupled with 100% effectiveness, ensures the maximum burst length to be fully transmitted within the olivary network. Intermediate current amplitudes modulate the number of spikes within the burst spikelet that are transmitted.

We considered an IO network consisting of 200 biophysically modelled cells embedded in a lattice arrangement (Fig 1B) [17]. Each IO dendrite was electrically coupled to 4 dendrites from nearby neighbour cells [17] via anisotropic gap junctions (i.e., directional electrical coupling, [7]) that could vary between 0 to 100%. We first tested the burst propagation between a pair of electrically coupled IO cells. The protocol involved the injection of a positive step current into one IO neuron (the left cell in Fig 1C), and the recording of the membrane potentials from both cells. A 100% coupling ensured a complete burst transmission from one IO cell to the other within 1-2 ms (Fig 1C). Second, we studied burst propagation across the entire network of 200 IO cells (Fig 1D). An excitatory step current was synchronously injected into a subset of central IO neurons, and we studied burst propagation as a function of the input current amplitude, whilst fixing the electrical coupling at 100%. We measured the cumulative distribution functions relative to the propagation of each burst spike, and we found a normal distribution with first spike: (μ, σ) ≈ (4 nA, 0); second spike: (μ, σ) ≈ (5 nA, 0.1); third spike: (μ, σ) ≈ (9 nA, 0.2); fourth spike: (μ, σ) ≈ (11.5 nA, 0.4) (Fig 1D). Therefore: an input current of 4 nA was sufficient to elicit 1 spike per burst across the entire 200 cell network; amplitudes larger than 13 nA elicited 4 spikes per burst (i.e., a complete burst propagation) across the entire network; and intermediate input amplitudes generated the propagation of intermediate IO spike burst lengths.

Then, we studied burst propagation across the IO network as a function of the electrical coupling strength. We considered an inhibitory input to the entire IO network to modulate the electrical coupling from 0% to 100% (Fig 2A). Given a fixed excitatory input current (15 nA), the strength of the electrical coupling significantly influenced the spike burst transmission across the network. Full burst propagation was guaranteed by a coupling strength higher than 85%, whereas, only 3 out of 4 spikes were propagated with, for instance, a 40% coupling (Fig 2B). The coupling strength also influenced the burst propagation time: a 100% coupling allowed for a full burst propagation through the entire IO network within 14 ms, whereas progressively lower levels of electrical coupling hindered the timing of burst propagation along with the number of spikes propagated (Fig 2C).

**Fig 2.**
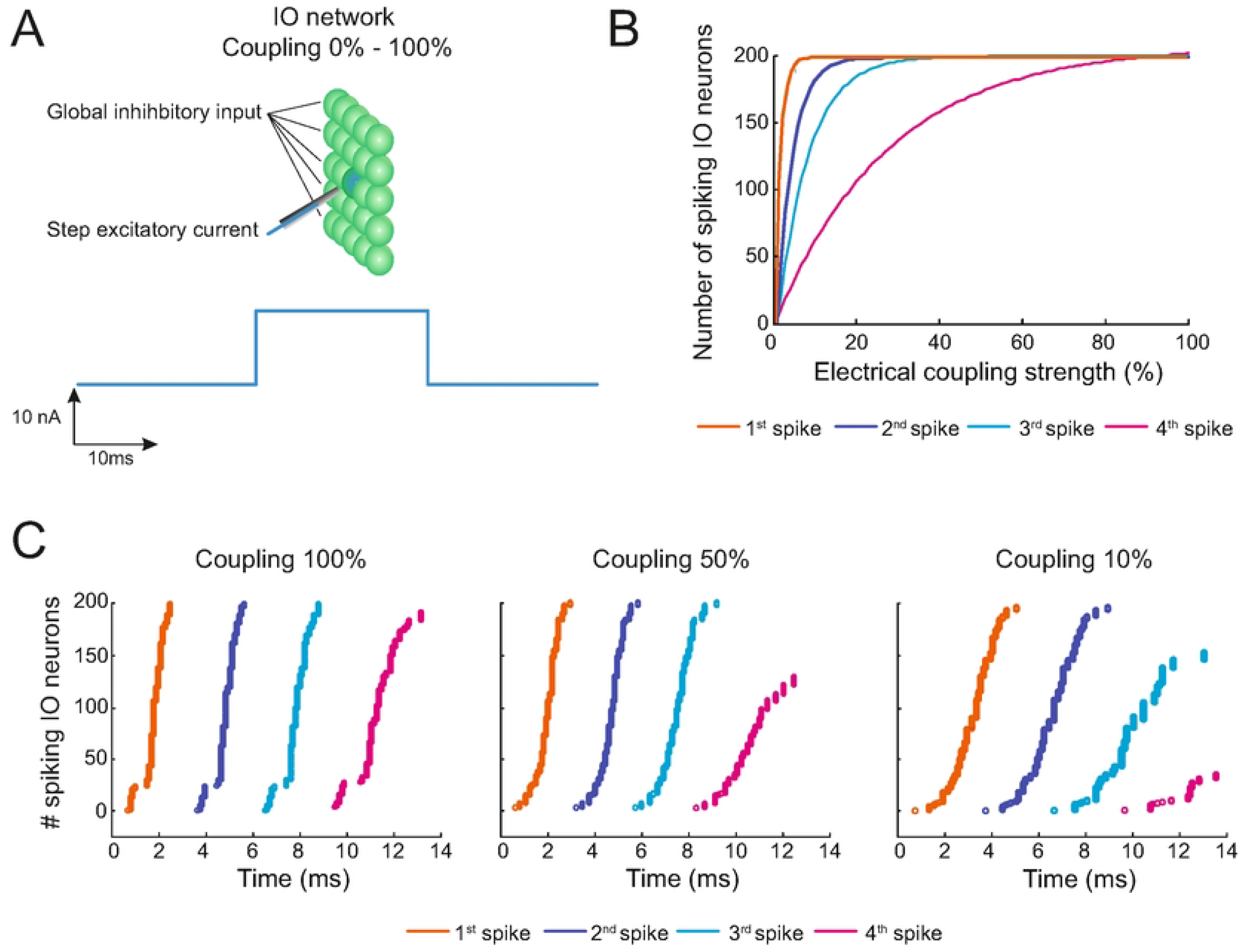
IO burst propagation properties via electrical coupling. **(A)** The schematic illustrates the simulation protocol, wherein a 10nA depolarising step current is applied for 20ms to the dendritic compartment of the central IO neuron in all subsets of 5×5 neurons within the lattice olivary network synchronously. The effectiveness of coupling is modulated to control burst propagation across the entire network. **(B)** Effective coupling varies from 0 to 100%, whilst observing IO bursts throughout the entire network. A 100% effective coupling ensures that the burst length is propagated within the lattice arrangement, whereas a 0% effective coupling leads to singular IO neural activations. Intermediate levels of effective coupling values act as regulators, influencing the transmission of burst length within the olivary network. **(C)** A 100% effective coupling enables the entire burst to propagate within 14 milliseconds. Reducing the coupling to 50% partially affects propagation, starting from the 4th spike within the spikelet and spanning the entire network. With a 10% effective coupling, transmission is affected from the 3rd spike within the spikelet across the entire network.

#### 2. IO subthreshold oscillations (STOs) and transmission of excitatory dendrite inputs

The IO neuronal model reproduced the subthreshold oscillation (STO) inner dynamics of real olivary cells [19-21], with a frequency of ∼10 Hz and an amplitude of ∼ 20 mV (Fig 3A). In the model (see Methods), during the hyperpolarisation phase, the somatic current I_k_ mediated by the K^+^ slow component channel dominated the initial rising of the IO membrane potential (red part in Fig 3A). From there, the somatic current I_ca_ (in blue), mediated by the calcium low threshold channel, a Ca^2+^-dependent K^+^ channel, further increased the membrane potential driving the hypopolarisation phase. Hereafter, either the STO continued (first circle) or a spike was generated (second circle), mediated through the I_Na_ and I_K_ currents at the olivary axon (green and purple lines, respectively). Then, the STO enters into its repolarisation phase thus resuming the oscillation cycle (Fig 3A). At the level of the IO network, our results confirmed that the electrical coupling amongst IO cells is essential to the synchronicity of STOs, but not their overall frequency (Fig 3B) [7].

**Fig 3.**
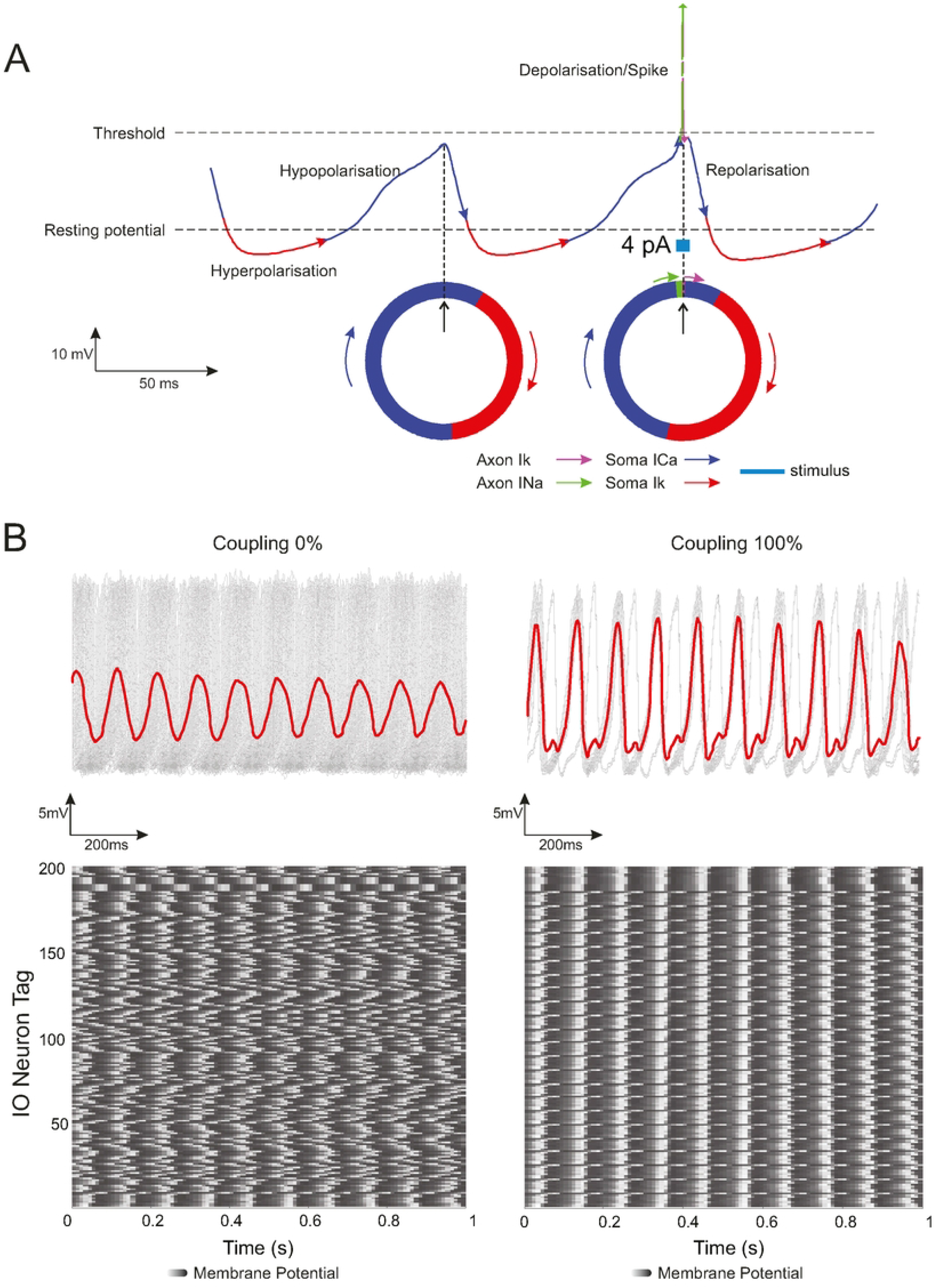
IO Subthreshold oscillations (STOs) within the olivary lattice arrangement and the impact of electrical coupling on the STO phase. **(A)** During hyperpolarisation, somatic current Ik (slow K+ channel) initiates the rise in IO membrane potential (in red). Subsequently, somatic current Ica (in blue) (low-threshold calcium channel and Ca2+-dependent K+ channel) enhances the depolarisation phase. This leads to either continued subthreshold oscillation (STO) or spike generation, mediated by INa and IK currents in the olivary axon (green and purple lines). The STO then transitions into repolarisation, restarting the oscillatory cycle. **(B)** STO frequency is shown for both electrically uncoupled (reducing coupling to 0%) and fully coupled (100% coupling) olivary network, revealing that overall frequency remains constant, with changes observed only in oscillation synchronicity. Coupling amongst IO neurons in the lattice arrangement transforms non-synchronous into synchronous oscillations. In the upper panels, the mean voltage of the coupled and uncoupled 200 IO network is depicted, whilst the lower panels provide a top-down view of the overall STO membrane potential of the IO network.

We studied the modulation of the STO phase by stimulating the centre of the IO network by a sequence of two excitatory inputs (Fig 4A). We sought to understand to what extent the relative timing of the two inputs (i.e., the interstimulus interval, ISI) would modulate (or possibly reset) the phase of IO STOs. We found that if the two inputs were delivered during different STO phases, they would cause either IO phase advances or delays (Fig 4B). When the second stimulus arrived during a hyperpolarisation period, it caused a delay in the STO phase between (0 – π). If the second stimulus arrived during hypopolarisation-depolarisation, it had the opposite effect, causing an advance in the phase between (π – 3π/2). Finally, when the second stimulus occurred during repolarisation, it caused again a delay in the phase between (3π/2 - 2π) (Fig 4B). Hence, a poor modulation or a reset of the STO phase could either partially or totally block the IO burst response to a sequence of excitatory synaptic inputs. The STOs thus provided a time-window gateway for the transmission of excitatory dendrite inputs occurring during either the IO hypopolarisation or the depolarisation time period.

**Fig 4.**
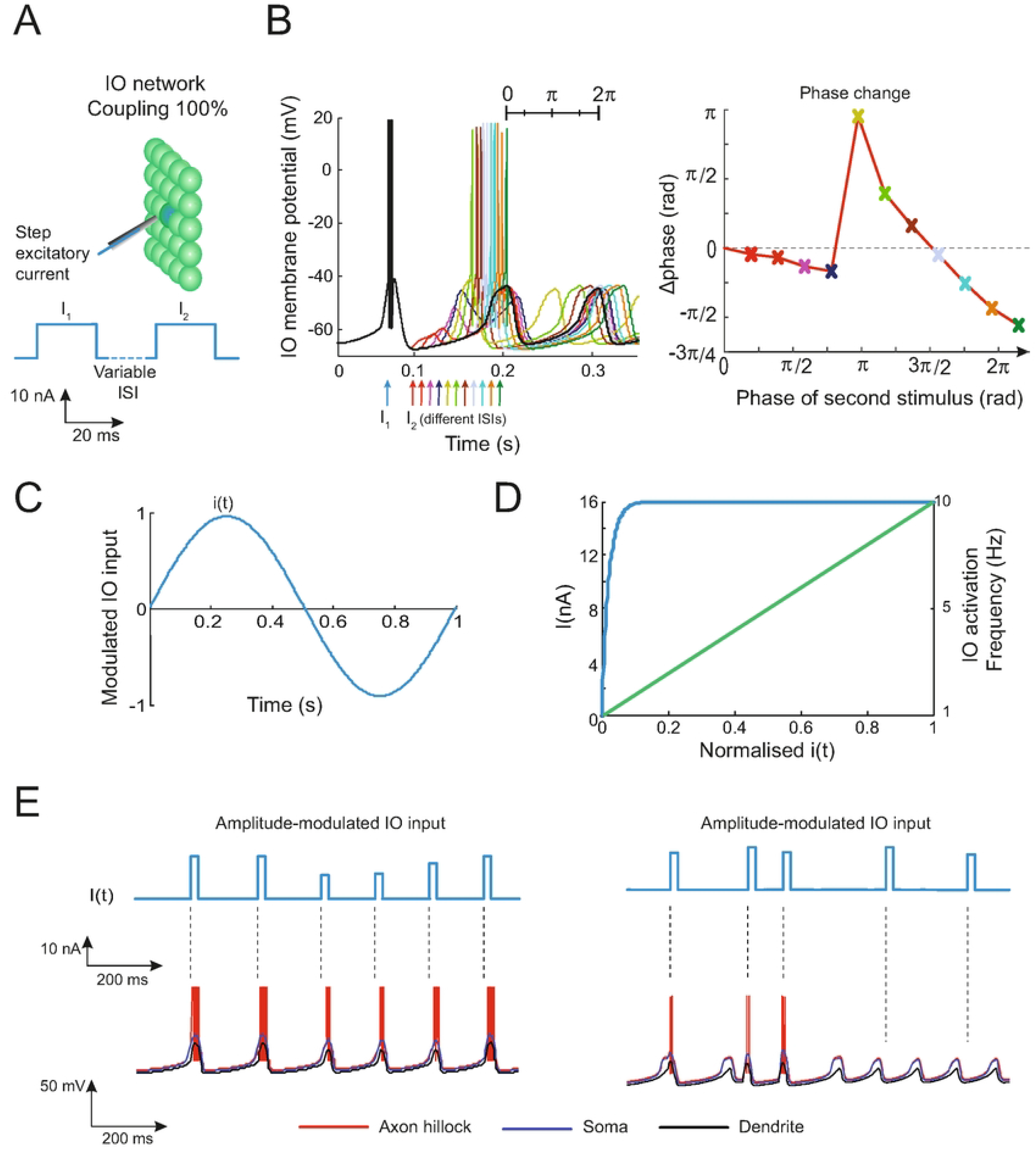
Modulation of subthreshold oscillation phase through sequential input instructive signals. **(A)** The schematic outlines the simulation protocol, involving the synchronous stimulation of the central IO neuron in all subsets of 5×5 neurons within the lattice olivary network. This stimulation maintains 100% effective coupling and is achieved using a depolarising current composed of a sequence of two stimuli, each with an amplitude of 14 nA during a 20 ms duration. Phase modulation was explored by varying the inter-stimulus interval (ISI) within the step sequence of the input instructive signal. **(B)** The left-hand side plot depicts the average membrane potential at the axon hillock of IO neurons within the lattice was plotted whilst varying the ISI [0 - 2π]. The right-hand side plot depicts STO phases in the olivary network responding to a variable ISI [0 - 2π] input signal. An early inter-stimulus occurring during the IO hyperpolarisation phase resulted in a phase delay within the range of (0 - π). Conversely, a late inter-stimulus occurring during the hyperpolarisation-depolarisation phase led to a phase advance within the range of (π - 3π/2). Finally, an inter-stimulus occurring during the repolarisation phase caused another phase delay within the range of (3π/2 - 2π). **(C)** Sinusoidal input curve mimicking retinal slip during r-VOR adaptation. **(D)** The Input current (depicted in blue) - Frequency (shown in green) IO Curve is used to generate temporal sequences of depolarising step current input stimuli for encoding retinal slip during VOR adaptation. The external input activity representing retinal slip sampling by IO activations follows a probabilistic Poisson process. In this process, the central IO neuron in all subsets of 5×5 neurons within the lattice olivary network is activated in the range of [1 - 10 Hz], where 1 and 10 Hz correspond to the minimum and maximum retinal slip values, respectively. Based on the normalised retinal slip signal i(t) (on the x-axis) and a random number η(t) ranging from 0 to 1, the central IO neuron in all subsets of 5×5 neurons within the lattice olivary network receives a depolarising step current. The amplitude of this current depends on the actual retinal slip amplitude when i(t) > η(t) i.e., the larger the retinal slip, the greater the amplitude of the input depolarising step current. **(E)** The upper panels illustrate two sets of temporal sequences of depolarising step currents used to encode the sinusoidal curve that simulates the retinal slip during r-VOR adaptation. The lower panels display the temporal evolution of voltage at the axon hillock, soma, and dendrite of the central IO neuron within a subset of 5×5 neurons. On the left-hand side, IO STO phase-locking is activated, meaning that retinal slip signalling is aligned with the IO hyperpolarization-depolarization STO phase, resulting in more precise sampling of the sinusoidal curve, i.e., no depolarising step current is lost. In contrast, on the right-hand side, an IO STO phase-free modulation is shown, where retinal slip signalling can occur at any time. In this case, several depolarising step currents are lost, and the spike burst lengths are diminished.

So far, only excitatory inputs with a fixed amplitude were considered. We therefore sought to study how amplitude-modulated inputs were transmitted by the IO network as a function of STOs’ phase. Again, we delivered sequences of excitatory inputs onto the centre of the IO network. However, the amplitude of these inputs was modulated according to a sinusoidal curve (Fig 4C). The injected step current was taken according to a probabilistic Poisson process, by comparing the sinusoidal function *i(t)* with a random number *η(t)* between 0 and 1. A positive input step current was injected at the centre of the IO network when *i(t) > η(t)*. The amplitude of the step current increased with the instantaneous |*i(t)*| value (Fig 4D), whilst its length remained fixed. When the step stimuli were well-timed with the hypopolarisation-depolarisation phases (STO phase locking with respect to the temporal input), the IO was able to properly encode and transmit the graded afferent signal properly (i.e., the length of IO burst responses reflected the amplitude of the input) (Fig 4E left). This was not the case in the absence of STO phase locking (Fig 4E right). IO responses were constrained to be below 10 bursts per second, consistently with those observed in neurophysiological recordings [22].

### B. Phase-locking of IO oscillations during r-VOR adaptation

#### 1. Cerebellar model for r-VOR adaptation

The IO network was integrated into a large-scale cerebellar model to learn r-VOR through adaptive feed-forward control (Fig 5; see Methods). We simulated a 1 Hz sinusoidal head rotation protocol (i.e., within the natural head rotation range of 0.05-5 Hz, [23]). The cerebellar model had to learn to move the eyes contralaterally with respect to the head rotation in order to minimise retina slips errors (i.e., the difference between actual and target eye movements, Fig S1).

**Figure 5.**
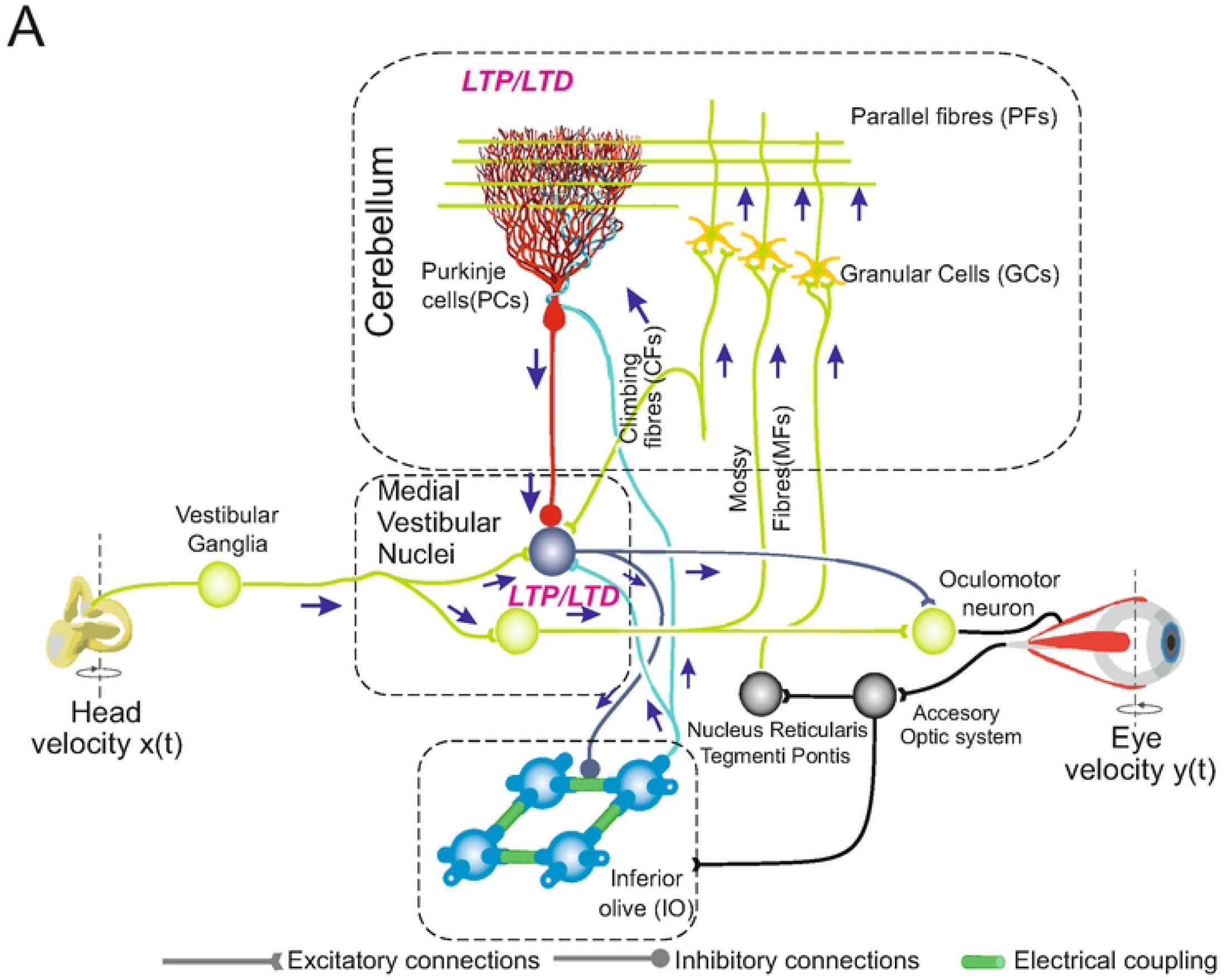
Cerebellum-dependent adaptation of vestibulo-ocular reflex (VOR). **(A)** Schematic representation of the main cerebellar layers, cells, and synaptic connections considered in the spiking cerebellar model. Mossy fibres (MFs) convey vestibular information onto granule cells (GCs) and medial vestibular nuclei (MVN). GCs, in turn, project onto Purkinje cells (PCs) through parallel fibres (PFs). PCs also receive excitatory inputs from the inferior olivary (IO) system. IO cells are electrically coupled and regulated via MVN-IO inhibitory connections. They deliver an instructive signal, which is the retinal slip, through the climbing fibres (CFs). Each MVN is inhibited by a PC and excited by an IO, both located at the same parasagittal band. MVN provides for the cerebellar output that ultimately drives oculomotor neurons. Spike-dependent plasticity occurs at PF-PC and MF-MVN synapses.

During head rotation, a population of 100 mossy fibres (MFs) encoded the primary vestibular inputs signalling head velocity to the cerebellar network. MFs projected excitatory afferents onto 200 medial vestibular nuclei (MVN) and 2000 granular cells (GCs). GCs expanded the coding space of MFs inputs [24] into 200 Purkinje cells (PCs) via parallel fibres (PFs, i.e., GCs’ axons). PCs were also driven by the climbing fibres (CFs, i.e., IO axons), which conveyed the teaching signal encoding retinal slip errors. The excitatory olivary CF collaterals along with inhibitory PC outputs contacted MVN neurons, which closed the loop through the MVN-IO inhibitory connections [25] conforming the olivo-cortico-nucleo-olivary (OCNO) loop [26]. MVN generated the cerebellar output that was sent to the oculomotor neurons, which ultimately drove eye movements. The OCNO subcircuit comprised two symmetric microcomplexes that compensated the ipsilateral head movement by controlling leftward and rightward eye rotations, respectively (see Methods). Cerebellar motor adaptation was driven by two spike-timing dependent plasticity (STDP) mechanisms at PF-PC and MF-MVN synapses. During 500 s of simulation, plasticity shaped PF-PC and MF-MVN synaptic efficacies (which were randomly initialised) to adapt VOR and reduce retinal slips [25, 27-29]

During r-VOR learning, the length of IO spike bursts (transmitted to target PCs via the CFs) had to encode the amplitude of retina image slips (i.e., errors). PCs’ complex spikes were linearly correlated with IO bursts (i.e., the spike number in Purkinje complex spikes depended linearly on the spike number in the CF bursts; [16, 25, 27, 28]. Hence, the different lengths of IO spike bursts could modulate the cerebellar adaptation capabilities, beyond an all-or-nothing learning paradigm [9].

#### 3. r-VOR adaptation requires IO STO phase-locking to error-related inputs

We tested the ability of the cerebellar model to perform r-VOR adaptation under two IO-dependent conditions: *(i)* in the presence of STO phase-locking to error-related inputs; *(ii)* in the absence of STO phase-locking, henceforth named as phase-free condition (i.e., with error signals arriving at any time with respect to IO STOs). STO phase-locking enabled a better time sampling of the error signal as well as a better encoding of its amplitude over time, which proved to be essential to mediate STDP at PF-PC synapses during r-VOR learning. As a consequence, the mean absolute error (MAE) (i.e., the difference between desired and actual contralateral eye movements) decreased over time, converging within 150 s (Fig 6A, red curve). Hence, IO STO phase-locking modulation allowed the cerebellum to maximise r-VOR accuracy, by optimising the r-VOR gain (i.e., the ratio between the antagonist eye and head displacements) and phase (Fig 6B; 1 Hz r-VOR gain = 1, phase = π), indicating that both eye position and velocity matched the ideal counter head movements (Fig 6C). By contrast, the r-VOR accuracy did not improve under the phase-free condition (Fig 6A, green curve), and neither the r-VOR gain nor the phase were optimised during learning (Figs 6B, C).

**Fig 6.**
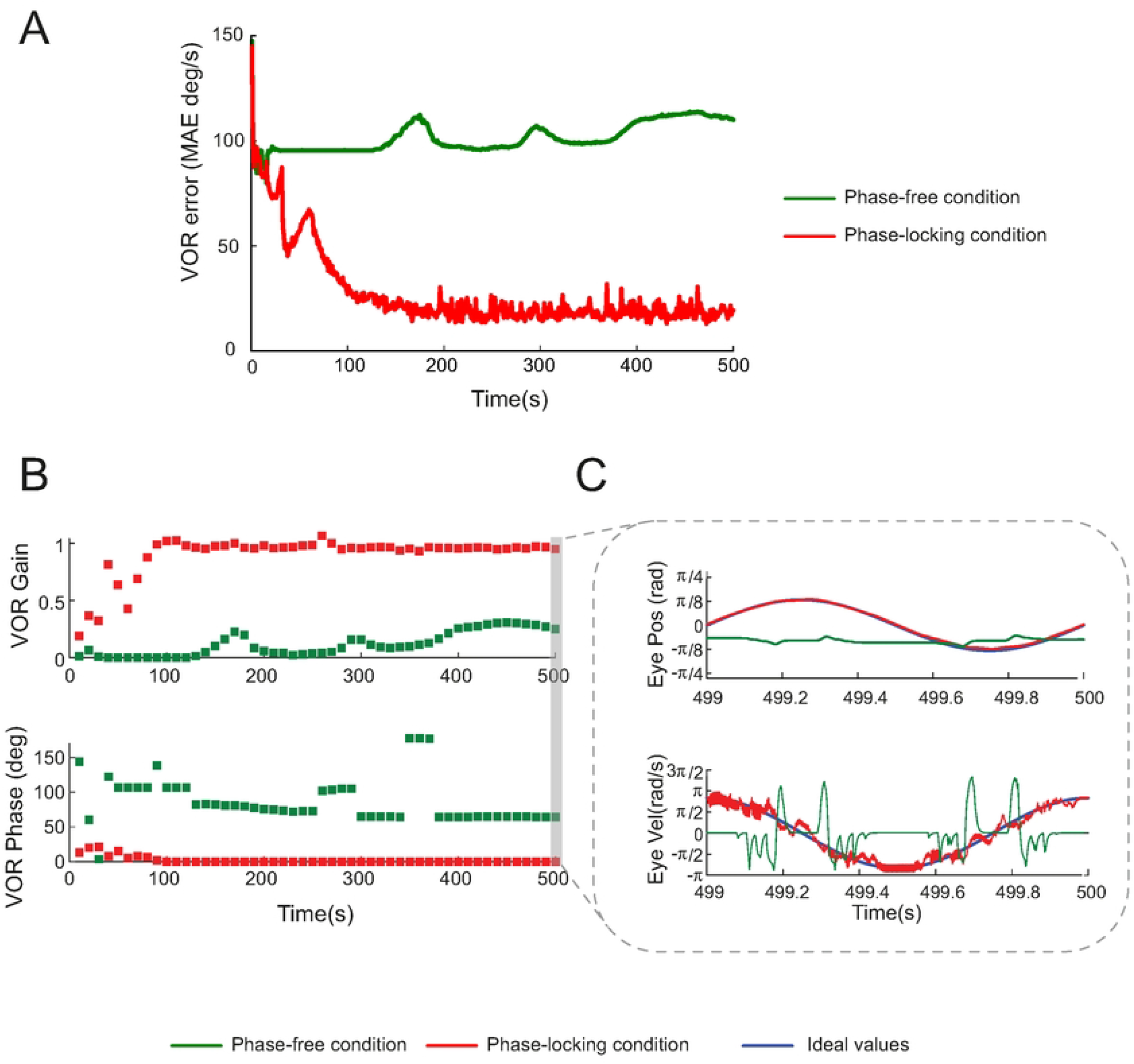
IO STO phase-locking vs phase-free modulation during rotational VOR acquisition. The olivary network in a lattice arrangement was integrated into a cerebellar network within a cerebellum-dependent feed-forward control scheme. This scheme was tested by assessing cerebellum-dependent r-VOR adaptation using a 1 Hz sinusoidal head rotation protocol during 500 seconds of simulation. Plasticity shaped PF-PC and MF-MVN synaptic efficacies to adapt rotational-VOR (r-VOR) gain and phase and reduce retinal slips. **(A)** MAE evolution for IO STO phase-locking and IO STO phase-free modulation during r-VOR adaptation (error = desired - actual sinusoidal r-VOR curve).**(B)** r-VOR gain and phase with IO STO phase-locking or phase-free modulation (A) r-VOR gain and phase evolution during r-VOR adaptation. (C) Actual and desired eye position and velocity at the end of the r-VOR adaptation process.

We found that the STO phase-locking condition elicited 5 times more IO bursts than the phase-free condition across r-VOR learning (Fig 7A). Also, cross-correlation analyses (i.e., between the spikes of IO burst responses during learning and r-VOR MAE values) suggested that the presence of STO phase-locking allowed the IO network to use the 4^th^ to 6^th^ spike of the bursts to grade the amplitude of the teaching signal driving STDP at PF-PC synapses (Fig 7B). The 1^st^ to 3^rd^ spike of the bursts were instead used to merely signal the presence of retina slip errors (the correlation between the 1^st^ to 3^rd^ spike burst and r-VOR MAE was constant; Fig 7B). Conversely, in the phase-free condition all spikes within IO bursts were equally correlated with the MAE error curve, thus indicating that bursts were only signalling errors (binary IO coding) without grading the teaching signal. In addition, during r-VOR learning the STO phase-locking condition generated a larger number of neural states of the IO network, whilst maintaining a small diversity of IO states (Fig 7C).

**Fig 7.**
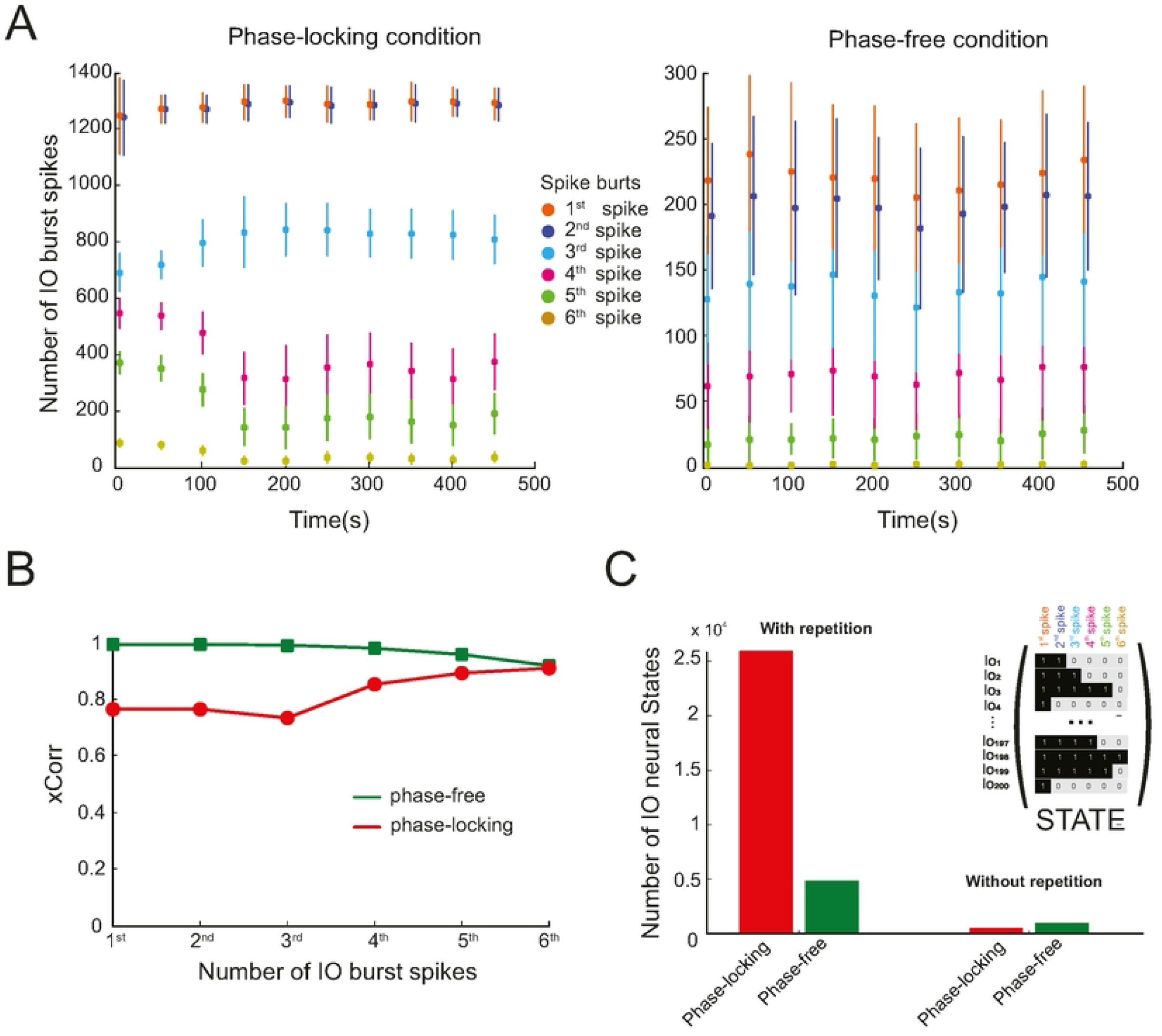
Spike bursts under IO STO phase-locking vs. phase-free modulation during r-VOR Acquisition. **(A)** IO burst number and length, i.e., spikes within the burst, under IO STO phase-locking and IO STO phase-free modulation during r-VOR adaptation. The IO olivary network elicited bursts to signal the presence or absence of retinal slips. STO phase-locking modulation allowed signalling the presence of retinal slips with four times more bursts under the same r-VOR protocol. Burst numbers remained consistent during r-VOR adaptation in both scenarios; however, IO STO phase-locking modified burst lengths along with r-VOR adaptation, i.e., retinal slip amplitudes were progressively declining. **(B)** Cross-Correlation analysis of IO STO phase-locking and phase-free modulations with their corresponding MAE curves obtained during r-VOR Adaptation (Fig 6A). When STO phase-locking modulation is used, the 3rd to the 6th spike within the burst grade retinal slip amplitude, i.e., correlation increases, whereas the 1st to 2nd spikes within the burst are used for signalling retinal slip only. In contrast, IO STO phase-free modulations employ the entire burst for signalling only. Note that IO STO phase-free modulations cannot achieve r-VOR adaptation (Fig 6A MAE curve), which means that only retinal slip signalling at any time is reflected in the burst length. **(C)** Neural states produced during r-VOR adaptation for IO STO phase-locking and IO STO phase-free modulation. Each neural state represents a binary matrix (200 x 6) that encodes the state of the olivary network whenever a burst is triggered. During r-VOR adaptation, it is observed that phase-locking modulation generates a larger number of unique neural states compared to phase-free modulation, albeit with slightly less repetition. On the other hand, phase-free modulation results in a somewhat lower repetition of neural states, primarily due to the random nature of the modulation, which involves free sampling.

#### 4. STO phase-locking & IO graded error coding improve r-VOR learning stability

We then comparatively analysed r-VOR adaptation under all-or-nothing versus variable error signalling. We ran a series of r-VOR learning simulations with IO STO phase-locking but under two conditions. In the all-or-nothing condition, we fixed the amplitude of the error signals received by the IO network (i.e., from minimum to maximum values, by increments of 25% of the range). Therefore, for a given simulation under the all-or-nothing condition, the IO network could only receive either a zero input or an input of a fixed amplitude. In the control variable error condition, we let the input to the IO network be modulated by the actual amount of retina slip (as in the previous r-VOR learning simulations).

We found that an all-or-nothing IO teaching signal up to the 75% of the maximum amplitude value did not allow the cerebellar network to minimise the r-VOR MAE over learning (Fig 8A, grey curves). Strikingly, when the all-or-nothing teaching signal was taken at the maximum value (100% of the range), the VOR MAE converged very rapidly (within less than 100 ms) to the optimal value (Fig 8A, black curve). However, VOR accuracy was not sustained over time, and the MAE began to slightly increase around 150 ms. By contrast, we found that a variable IO teaching signal amplitude allowed for both VOR MAE minimisation (convergence within about 150 ms) and learning stability (Fig 8A, red curve). These results were reflected in the evolution of the r-VOR gain and phase across learning (Figs. 8 B and C, respectively). An analysis of the eyes position and velocity curves at the end of learning (i.e., at 500 s) confirmed a better match, with respect to the ideal profiles, in the presence of a variable teaching signal amplitude as compared to an all-or-nothing one (Figs. 8D, E). Finally, whilst assessing the factors beneath the better r-VOR learning performance provided by a variable teaching signal, we found that this condition allowed a larger number of IO neural states to be generated, as compared to binary error signalling (Fig 8F). Interestingly, a larger number of IO neural states led to no FFT harmonics in eye position and velocity curves (Fig 8G), resulting in better r-VOR learning stability.

**Fig 8.**
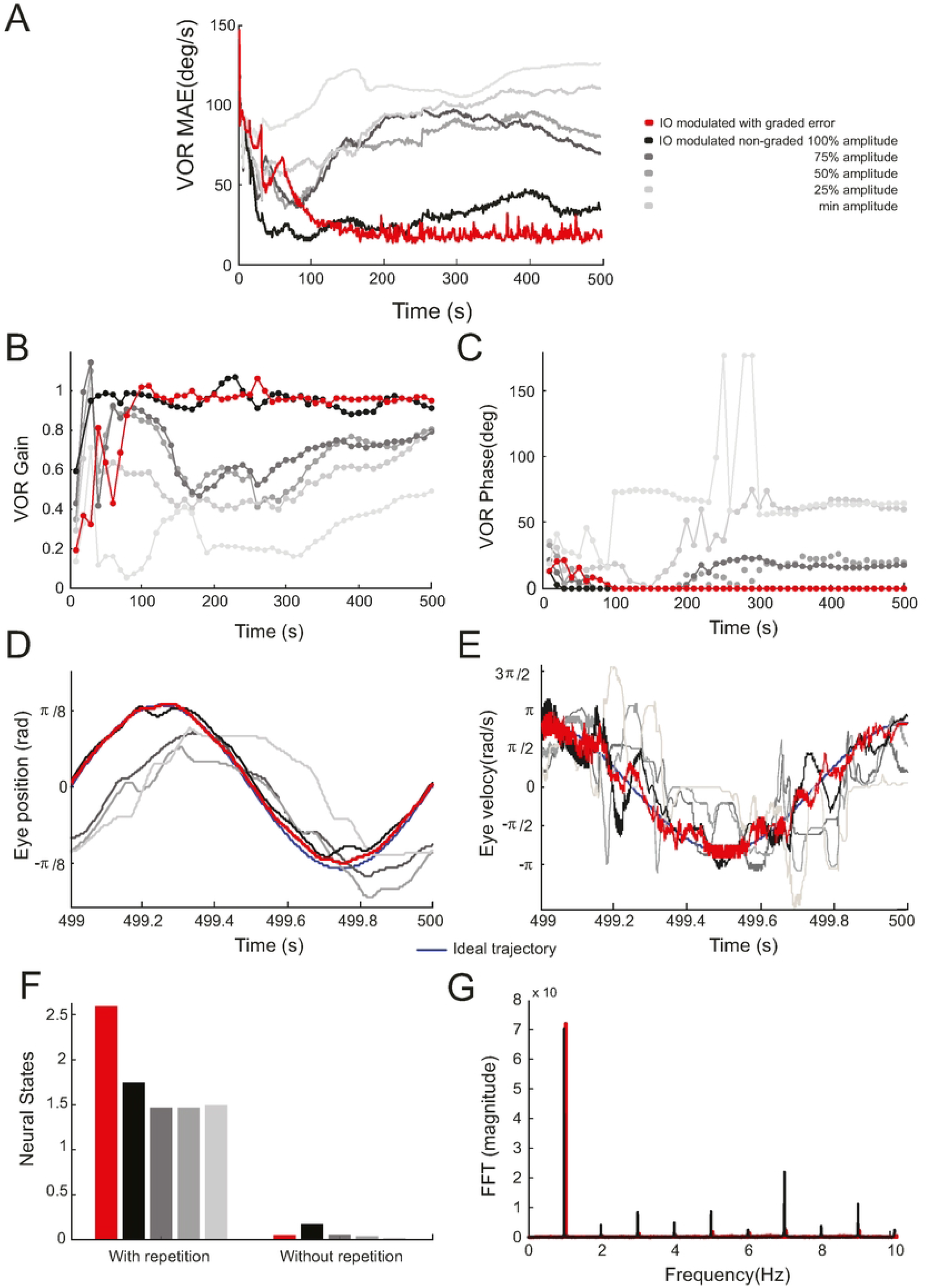
IO STO phase-locking behaviour under burst length modulation. We build upon the experimental setup introduced in Fig 7, focusing exclusively on the modulation of IO STO phase-locking. Specifically, we examine the impact of different retinal slip amplitudes, comparing fixed burst lengths whilst controlling retinal slip amplitude in 25% increments. We also consider variable burst lengths regulated through retinal slip amplitude modulation. **(A)** Evolution of MAE during r-VOR adaptation. The MAE serves as a measure of how closely our model aligns with the desired sinusoidal r-VOR curve. Notably, we find that the graded instructive signal configuration (represented by the red curve) results in a further decrease in MAE and better stability. **(B-C)** Evolution of r-VOR Gain and Phase during r-VOR adaptation. These measurements show similar performance between graded and non-graded instructive signal amplitudes when retinal slip amplitude is fixed at its maximum (100%). **(D-E)** Display of actual and desired eye position and velocity at the end of the r-VOR adaptation process assessing how well our setups match the desired outcome. **(F)** Analysis of Neural States during r-VOR Adaptation. The graded instructive signal amplitude (retinal slip) generates a greater number of neural states with less variability compared to the non-graded instructive signal amplitude when the retinal slip amplitude is fixed at its maximum. Notably, an initial rapid decrease in MAE for the non-graded instructive signal configuration (see Fig A) suggests that neural state variability aids in r-VOR convergence. **(G)** Fast Fourier Transforms (FFT) of eye velocity. In this analysis, it is compared the FFT of eye velocity between graded and non-graded retinal slip amplitude (fixed at 100%). The FFT of the non-graded instructive signal amplitude (fixed at 100%) reveals larger odd and even harmonics, indicating a poorer fit to the ideal r-VOR curve, despite gain and phase measurements suggesting optimal performance. Note that gain and phase measurements only consider the first harmonic (see methods).

#### 5. STO phase-locking & GABAergic regulation of IO electrical coupling improve r-VOR adaptation

In all previous r-VOR simulations, we did not activate the GABAergic MVN-IO projections in the model (Fig 5). Here, we considered them in order to account for their known role in modulating the electrical coupling amongst IO cells [30]. We sought to investigate to what extent the modulation provided by these MVN-IO inhibitory synapses could play a role in increasing IO neural state diversity (on top of the larger number of neural states provided by variable teaching signalling, shown in Fig 8F).

We ran a series of r-VOR simulations to compare two adaptation scenarios: a condition with MVN-IO inhibitory regulation of IO coupling, and a condition without it. In both conditions, we preserved the IO STO phase-locking to error-related inputs. For the condition “with MVN-IO inhibition”, we first ran a sensitivity analysis to set the MVN-IO synaptic weights in order to optimise the VOR MAE function. Then, for each condition, we simulated 100 r-VOR adaptation experiments (again based on a 1 Hz sinusoidal head rotation during 500 s). We found that whilst the total number of IO neural states diminished, an increase of state diversity was associated with the presence of MVN-IO GABAergic modulation (Fig 9A), which resulted in a significantly better VOR accuracy (Fig 9B). Hence, even if MVN-IO inhibition was not a necessary condition for r-VOR adaptation, it contributed to facilitating r-VOR learning.

**Fig 9.**
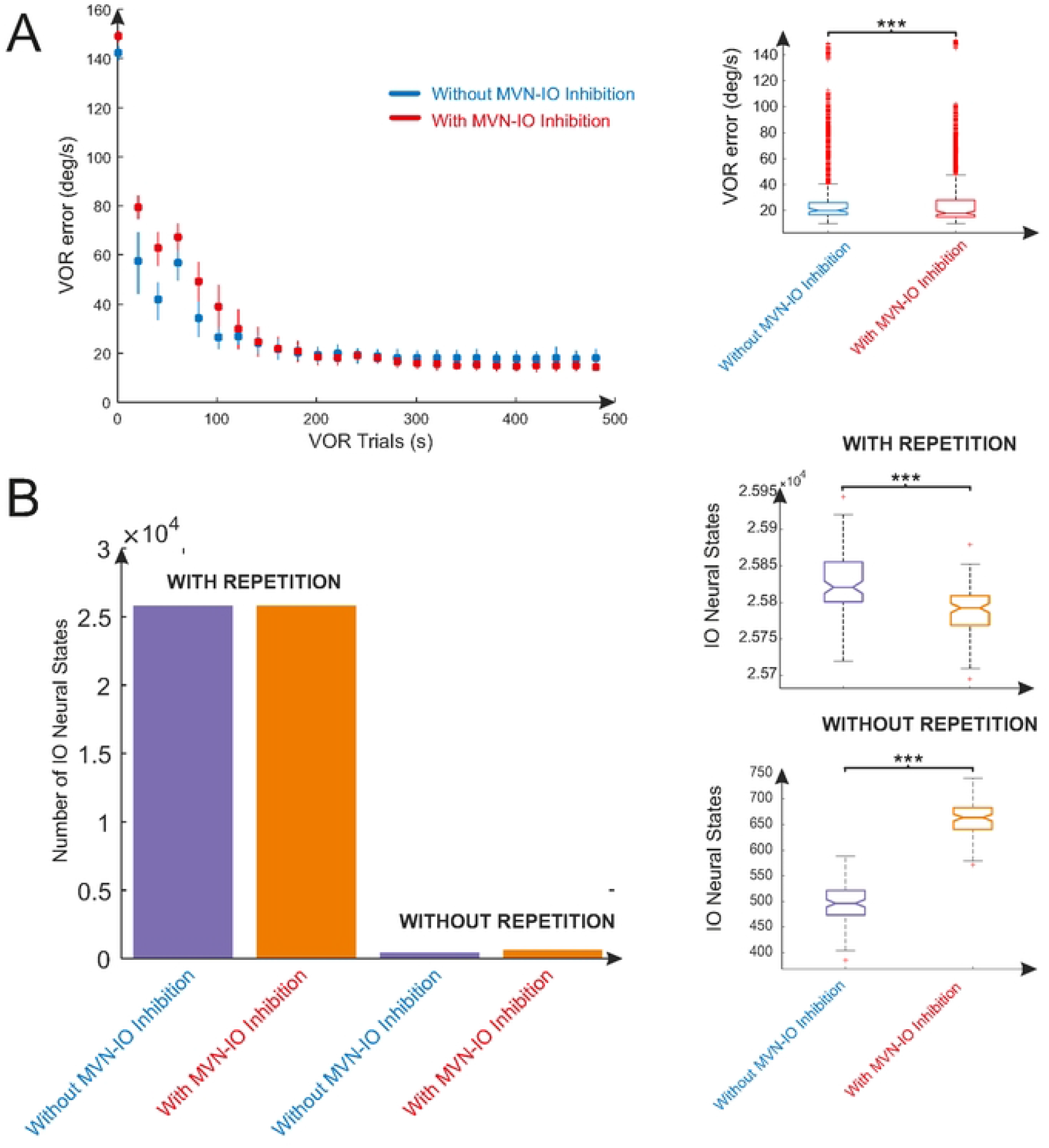
Enhancing r-VOR accuracy by regulating IO-IO coupling through MVN-IO inhibitory afferents. We maintained the experimental setup from Fig 9, where variable burst lengths were regulated solely through retinal slip amplitude modulation. In this configuration, we enabled/disabled MVN-IO inhibitory connections. **(A)** Evolution of MAE during r-VOR adaptation (mean and standard deviation each 10 seconds). MAE measures the deviation between the desired and the actual sinusoidal r-VOR curve. An ANOVA statistical test confirms significant differences in MAE with and without MVN-IO inhibitory connections, indicating a lower MAE and, consequently, a more accurate r-VOR adaptation in the presence of EC regulation via MVN-IO afferents. **(B)** Neural States generated with and without MVN-IO inhibitory connections. A statistical test (ANOVA) confirms significant differences in the neural states generated. The presence of EC regulation via MVN-IO afferents results in a lower overall number of neural states but with increased diversity, leading to a more accurate r-VOR adaptation.

## III. Discussion

The olivary nucleus plays a crucial role in cerebellar adaptation by influencing synaptic plasticity between parallel fibres and Purkinje cells. This study suggests that the dynamics of the inferior olive (IO) network, particularly the phase of subthreshold oscillations (STOs) in response to excitatory inputs, is necessary for sensorimotor adaptation. We created a model of IO cells to mimic real activity and we integrated it into a cerebellar model to study vestibular ocular reflex (VOR) adaptation. Our results confirmed that (i) the presence of STOs generated an opportunity modulation time window occurring during the IO hipopolarisation-depolarisation time period; (ii) STO phase-locked modulation during this period allowed the retinal slip [31] signals at low frequencies [1 - 10Hz] to be adequately sensed and naturally graded via spike-burst lengths; (iii) this modulation together with electrical coupling (EC) allowed for the generation of enough olivary neural states to ensure VOR adaptation; (iv) a wider variety of neural states increased VOR adaptation converging speed. Neural state variety was found to be increased thanks to the EC regulation via the GABA nuclei projections onto the olivary network. These results allow us to postulate a theory on the olivary system operating as a burst-amplitude modulator (see below).

### A. IO operates as a master clock and as teaching signal during VOR adaptation

Three conditions were fulfilled by the presented IO system to act as a master clock [32]: *(i)* The membrane potential of the Hodgkin-Huxley neuronal model was operating as a continuous and accurate carrier frequency that acted as the reference signal. Our HH model membrane potential, acting as the reference signal, was able to generate STOs with a precise 10 Hz periodicity, thanks to the dynamic interactions of ionic channels. These STOs provided multiple modulation opportunities during their hypopolarisation-depolarisation phases, i.e., temporal windows. (ii) The natural range of the r-VOR is from 0.5 to 5 Hz, with our operating frequency being within 1 Hz. Since the temporal frequency of the sensed signal (1 Hz) is lower than the “master clock period” (10 Hz), this ensures sufficient temporal precision [33]. (iii) A timely sequence of external input activities reaching IO cells could control the initiation and termination of STOs, allowing for the correlation of IO timing signals with retinal slip signals (an instructive signal) that drive the specific operation of the r-VOR model system. Note that the external input activities, that is the retinal slip signals, resulted from a Poisson sampling of the retinal slip stimulus.

Our modelled olivary system was also able to convey a low-firing rate instructive signal which is typically based on retinal slip amplitude at approximately 1 to 10 Hz. This instructive signal helped the cerebellum compensate for head rotary movement by controlling and adapting the contralateral eye movements (r-VOR), despite the diminished signal transmission capability of the olivary system due to its low-frequency operation [30]. To ensure a proper representation of the entire retinal slip region over trials, i.e., desired vs. actual eye velocity, we generated external input activity driven towards the IO cells using a probabilistic spike sampling of the retinal slip signals (instructive signal generation) according to a Poisson process, whilst maintaining the IO activity between 1 and 10 Hz per fibre (similar to electrophysiological data [22]). This approach allowed us to accurately sample the retinal slip evolution even at such a low frequency, as supported by previous studies [25, 27-29, 34, 35]. We assumed that the CF triggered burst signals based on retinal slip magnitude, supported by two findings in awake mice: (i) CF-triggered signals gradually increases with the duration and pressure of periocular stimuli [36, 37] and (ii) The amplitude of CF-triggered signals onto PC dendrites is graded and represents information about the intensity of sensory stimuli [18, 38].

We found that the spike burst triggered by CF into the PC dendrites were neither fully binary nor fully graded. Interestingly, the 1^st^ and 2^nd^ spikes within the burst were used to indicate the binary presence of the retinal slip signals, whilst the 3^rd^ to 6^th^ spikes were used to naturally grade the retinal-slip signal amplitude. This CF spike burst modulation, according to the retinal slip amplitude, did not require the GABAergic nuclei cells to decrease the IO electrical coupling, which can cause the olivary network synchronicity to break [14]. Instead, we found that the GABAergic nuclei cells’ ability to desynchronise the olivary network played a role in providing more accurate r-VOR adaptation. We confirmed the GABAergic nuclei action increased the non-redundant neural states in the olivary network during r-VOR adaptation. This increase in the IO information transmission capability contributed to a more precise r-VOR adaptation.

### B. HH IO phase modulation and criticality of STOs

Our IO HH three compartment model, based on previous studies [7, 19], was designed to alleviate the computational load whilst maintaining the main morphological and functional properties of the olivary system, especially the generation of spiking bursts at the axon hillock. The PC HH single-compartment model was also able to reproduce the spiking modes of Purkinje cells, including tonic, pause, and burst firing patterns. The PC burst reflected a perfect burst transmission from its corresponding CF [27]. The IO ionic channel dynamics in our model caused STOs to naturally appear at 10Hz, generating opportunity modulation time windows for the IO spike burst responses when following the sensorial stimulation of the IO dendrites. We also found that the temporal input sequences towards the olivary system were able to start/reset the IO STOs generation, and the IO spike burst modulations only occurred properly during their hipopolarisation-depolarisation time periods, indicating STOs as a conditional complex spike gating mechanism [7, 8].

These two timing facts pointed to a phase-locked modulation of the STOs during the IO hipopolarisation-depolarisation period. The start/reset mechanism adjusted the IO voltage reference signal to the occurrence of the Poisson sampling of retinal slip signal, whilst the IO hyperpolarisation-depolarisation period adjusted the IO burst length modulation to the amplitude of the Poisson-sampled retinal slip signals during VOR adaptation.

### C. The purpose of the IO STOs: Olivary system operating as a burst-amplitude modulator (BAM), a theory

In the context of sensory neural processing, the necessity of modulation within the olivary nucleus becomes evident. When attempting to convey multiple sensory stimuli directly to the IOs and PCs downstream without modulation, an inherent issue arises. This issue stems from the fact that all sensory stimuli sharing the same frequency range would saturate the IO-PC-MVN neural circuitry. This is similar to attempting to tune into multiple radio stations operating on the same frequency simultaneously. As a result, the absence of olivary modulation only allows for the transmission of one sensory stimulus at any given time. To address this limitation, a modulation technique involving STOs could be used. The STO modulation shall effectively shift the frequencies of sensory stimuli to higher ranges, typically around 10 Hz. Furthermore, it shall enable the assignment of distinct frequencies to individual sensory stimuli, similar to the concept of amplitude modulation (AM) in radiofrequency transmission. However, our STO modulation does not operate over the amplitude of the carrier signal, represented by STOs amplitude, based on the sensory message. Instead, it varies the lengths of complex spike bursts, ensuring a diverse representation of the sensory input.

Interestingly, in amplitude modulation (AM), the carrier signal is modulated by the message signal through multiplication. Additionally, a constant value is added to the message signal. This dual action ensures that when the message signal is at its smallest values, the carrier signal effectively disappears, which is a technique to prevent over-modulation (Supplement S2). Over-modulation, in this context, is akin to the phase of the carrier signal reversing, which can lead to extreme distortion in the subsequent demodulated signal (Supplement S3). Similarly, within the context of IO STOs phase modulation acting as the carrier signal, the sensory stimulus, representing the message signal, is characterised by a variable amplitude input current. The amplitude of this current is equivalent to the magnitude of retinal slip amplitude. This dynamic input, when injected into the IO dendritic compartment, results in the generation of complex spike bursts of varying sizes. Furthermore, the first (1^st^) and second (2^nd^) spikes within the CS bursts function as a binary signal to denote the presence or absence of the stimulus. Note that these 1^st^ and 2^nd^ spikes also carry a constant modulation, which shall contribute to the effective reduction of the carrier signal amplitude, essentially causing the STOs to vanish at the lowest sensorial stimulus values, thereby preventing over-modulation.

Significant parallels exist between AM demodulation and the mechanisms occurring in the MVN, specifically, the cerebellar neural decoding process. MVN neurons are recognised for their ability to encode various frequency oscillations related to horizontal linear motion. Notably, the medial section of the MVN has been observed to respond to low-frequency stimulation, typically in the range of 0.5 to 1.0 Hz in studies involving rats [39]. Further investigations conducted in vitro have identified a distinct subtype of neurons within the vestibular nuclei, labelled as ‘type B’ that exhibit a form of adaptation in their firing rate in response to depolarising current steps. This adaptive behaviour displays resonance at frequencies within a range relevant to behaviour, facilitating synchronisation with the peaks of incoming stimuli [40, 41]. Additionally, modelling of the vestibular nuclei suggests the presence of membrane potential oscillations in response to step current inputs, which is indicative of phenomena that might manifest in vivo [42]. Given these observations, it is plausible to consider the oscillations within MVN as a vital aspect of cerebellar decoding. In essence, they can be viewed as a form of a product detector demodulator [43]. The cerebellum demodulation process shall combine the modulated sensorial stimulus with input from inhibitory PC afferents and vestibular signals from MF afferents, incorporating a local oscillator represented by type B vestibular nuclei neurons. Crucially, these type B vestibular nuclei neurons must oscillate at the same frequency as IO STOs, effectively acting as the carrier signal (Supplement S3). The output from MVN shall carry several robust cerebellar outputs, and it shall contain a signal within the frequency range of the sensory stimulus (message). This, in turn, shall lead to the faithful reproduction of the original modulating signal, which represents the sensory stimulus after Spike-Timing-Dependent Plasticity (STDP) learning adaptation (Fig 10).

**Fig 10.**
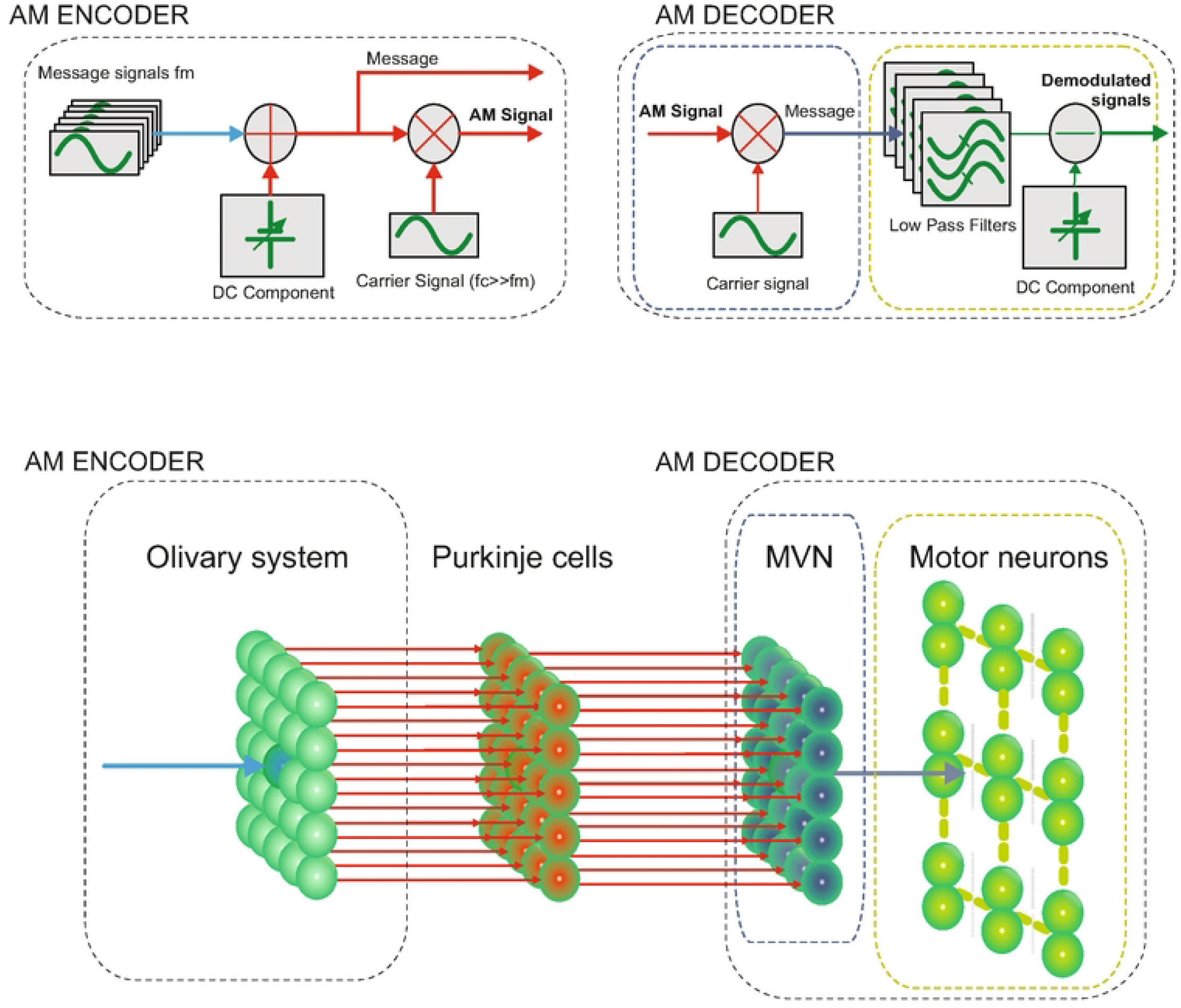
Analogies between amplitude modulation in radio transmission and burst-amplitude modulation in the olivary system. The olivary network STOs use a phase-locked modulation in amplitude. However, the STOs modulation varied the burst lengths according to the retinal slip amplitudes instead of varying the amplitude of the carrier signal. Consequently, all the sensorial stimuli sharing the same frequency range could simultaneously be transmitted. The olivary system may play the same role as an AM encoder during the input stimuli transmission to downstream cerebellar layers, whereas the MVN together with the motor neurons may play the same role as an AM decoder for stimuli reconstruction at the cerebellar output.

A product detector demodulator uses a direct conversion reception method to extract the message signal, which is the most straightforward approach for receiving information transmitted by a carrier (Supplement S3). An essential component for this process is a simple low-pass filter, known for its effective selectivity [44].The behaviour of motor ocular neurons aligns with the operation of a low-pass finite impulse filter (FIR), as observed in previous research [45]. This alignment may contribute to the process of demodulation in the medial vestibular nuclei (MVN) (Fig 10).

## IV. Materials & Methods

### A. VOR Analysis and Assessment

We simulated the horizontal VOR (h-VOR) during sinusoidal (∼1 Hz) whole-body rotations [46]. VOR gain was determined as the ratio between the first harmonic amplitudes of the eye and head velocity Fourier transforms:

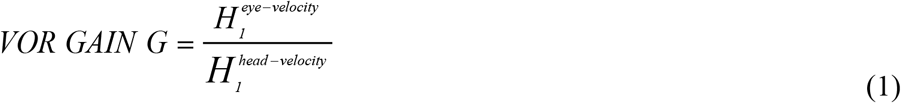

Conversely, VOR shift phase was calculated as the cross-correlation of the eye (e) and head (h) velocity time series:

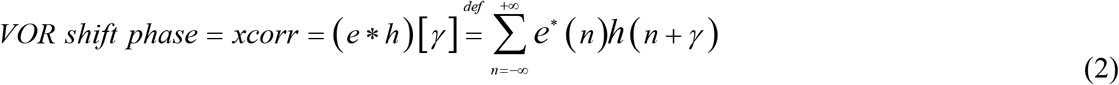

Here, e^*^ represents the complex conjugate of e, and γ the lag indicates the shift phase. After normalisation, the ideal eye and head velocity lag is ± 0.5, with cross-correlation values ranging from -1 to 1. This range is equivalent to a phase shift interval of [-360º, 360º].

### B. VOR Mechanical Circuitry

The cerebellum operates as a biological feed-forward controller within a control loop. Its output drives adaptation from the MVN through a series of motor neurons, nerve fibres, and muscles, ultimately affecting eye movement. We modelled this pathway using the EDLUT neural simulator [47-49] as VOR (Vestibulo-Ocular Reflex) mechanical circuitry defined by a continuous-time mathematical model:

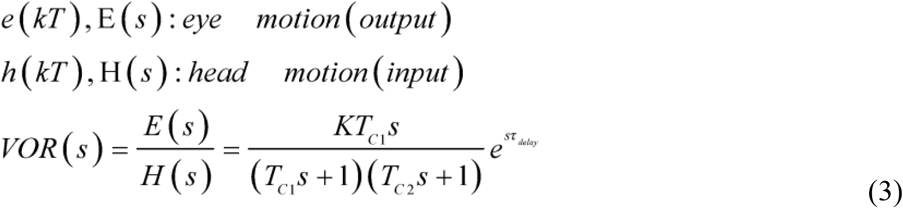

This model consists of four parameters: *Q = [K, T*_*C1*_, *T*_*C2*_, *τ*_*delay*_]. The delay parameter *τ*_*delay*_ accounts for the time it takes for signals from the inner ear to reach the brain and eyes, estimated to be around 5 ms based on the number of synapses involved in the VOR [50, 51]. It is also included in the cerebellar sensorimotor pathway delay (see the STDP section) [25, 27, 28]. The gain parameter *K* represents the inability of the eyes to perfectly track head movements and it is assumed to fall within the range of 0.6 to 1 [50, 51]. *T*_*C1*_ reflects the dynamics associated with the semicircular canals and additional neural processing. These canals act as high-pass filters, because after a subject has been put into rotational motion, the neural active membranes in the canals slowly relax back to resting position, so the canals stop sensing motion. Based on the mechanical characteristics of the canals, combined with additional neural processing which prolongs this time constant to improve the accuracy of the VOR, the *T*_*C1*_ parameter is estimated to be between 10 and 30 seconds [50, 51]. Finally, *T*_*C2*_ characterises the oculomotor plant dynamics, including the eye, muscles and attached tissues, with TC2 assumed to be between 0.005 and 0.05 seconds.

To obtain the temporal response for the VOR transfer function, we need to calculate the inverse Laplace transform, taking into account that the delay is modelled and inserted within the control loop (Eq 4).

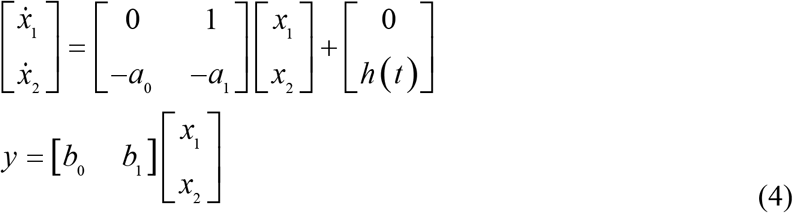

Where: *a*_0_ =1/(*T*_*c*1_*T*_*c*2_);*a*_1_=*a*_0_=(*T*_*c*1_+*T*_*c*2_)/ (*T*_*c*1_*T*_*c*2_);*b*_0_=0;*b*_*1*_=*KT*_*c*1_/(*T*_*c*1_*T*_*c*2_);The VOR plant model parameters were fine-tuned using a genetic algorithm to align with experimental and clinical observations1 [50-52]. The resulting parameter values are: *K* = 1.0, *T*_*C1*_ = 15, *T*_*C2*_ = 0.05.

### C. Cerebellar Spiking Neural Network Model

The cerebellar circuit, modelled as a feed-forward loop, effectively compensated for head movements through contralateral eye movements (see Fig 5). This cerebellar network comprised five neural populations: mossy fibres (MFs), granule cells (GCs), medial vestibular nuclei (MVN), Purkinje cells (PC), and inferior olive (IO) cells [53-57]. This cerebellar model was implemented in EDLUT [47-49], an open-source, spiking-based neural simulator designed for efficient computation and embodied experimentation.

#### Mossy fibres (MFs)

We modelled 100 MFs as input neurons responsible for transmitting sensory-motor information to both GCs and MVN. In line with the functional principles of VOR models for cerebellar control [58], MF activity ensembles were generated to follow a 1 Hz sinusoidal pattern, with a new MF ensemble for each 2 ms simulation step, encoding head [58-60]. The overall MF activity was organised into non-overlapping and equally sized neural subpopulations to maintain a consistent firing rate for the MF ensemble over time. Note that two different times corresponded to two different subgroups of active MFs, ensuring overall constant activity (see Network connectivity parameters summarised in Table 1).

**Table 1.** Cerebellar network topology parameters. (Dash lines indicate not applicable)

| Neurons |  | Synapses |  |  |  |
| --- | --- | --- | --- | --- | --- |
| Pre-synaptic neurons (number) | Post-synaptic neurons (number) | Number | Type | Initial weight(nS) | Weight range(nS) |
| 2000 GCs | 200 PCs | 400000 | AMPA | rand | [0,4] |
| 200 IO | 200 PCs | 200 | AMPA | 40 | – |
| 100 MFs | 2000 GCs | 8000 | AMPA | 0.35<sup>a</sup> | – |
| 100 MFs | 200 MVN | 20000 | AMPA | 10 | [0, 10] |
| 200 PCs | 200 MVN | 200 | GABA | 1.5 | – |
| 200 IO | 200 MVN | 200 | NMDA | 7 | – |
| 200 IO | 200 MVN | 200 | AMPA | 1 | – |
| 200 MVN | 200 IO | 200 | GABA | 0.15 | – |
| IO to IO configuration, there are 5x5 IO neuron squares arranged in a lattice pattern. These squares are connected radially, extending from the centre to each corner of the 5x5 square, resulting in a total of 200 IO neurons. |  | 320 | GAP<br>JUNCTION | 1 | – |

<sup>a</sup> We used specific parameters to generate offline GrC activity. This activity remained constant throughout the r-VOR adaptation process. This pre-generated GrC activity was preloaded during computation to expedite the simulation.

#### Granule cells (GCs)

The granular layer consisted of N = 2000 GCs (Leaky Integrate & Fire (LIF) neurons) and functioned as a state generator [61-64]. The inner dynamics of the granular layer produced time-evolving states comprising non-overlapping spatiotemporal patterns that were consistently activated in the same sequence during each learning trial (1 Hz rotation for 1 second). Despite receiving a constant MF input encoding each second of the 1 Hz learning trial, the granular layer generated 500 different states. Each state was composed of four non-recursively activated GCs [65].

#### Purkinje cells (PCs)

200 PCs were modelled using a single compartment Hodgkin-Huxley (HH) model with five ionic currents, allowing them to replicate the tri-modal spike modes (tonic, silence, and bursting) observed in PCs [27, 66].These PCs were divided into two subpopulations of 100 neurons each. Each subpopulation received inputs from 100 CFs arranged in a lattice configuration [17]. These CFs encoded the difference between both clockwise or counter clockwise eye and head movements. Additionally, each PC received 2000 PF inputs. Given that PCs are innervated by approximately 150,000 PFs [67], the weights of the PF-Purkinje cell synapses in the model were adjusted to match the biological excitatory drive. Each of the two subgroups of 100 Purkinje cells targeted 100 MVN cells through inhibitory projections. The MVN cells were responsible for generating either clockwise or counter clockwise compensatory motor actions, ultimately driving the activity of agonist/antagonist ocular muscles.

#### Inferior olive (IO)

200 IO cells were modelled using a three-compartment Hodgkin-Huxley (HH) model equipped with seven ionic currents and electrical coupling. This HH model accurately reproduced both the spike burst and the subthreshold oscillations observed in the IO [68]. The neural layer was divided into two subpopulations of 100 neurons each, arranged in a lattice configuration[17]. These two subpopulations were responsible for handling clockwise and counter clockwise sensed errors. CFs transmitted the instructive signal (retinal slips) from the IOs to the populations of PCs. Each CF made contact with one PC and one MVN cell. The external input activity of IO cells was generated using a probabilistic Poisson process. Based on the normalised retinal slip current curve i(t) and a random number η(t) ranging from 0 to 1, the central IO neuron in all subsets of 5×5 neurons within the lattice olivary network received a depolarising step current (see Fig 4A).The amplitude of this current depended on the actual retinal slip amplitude when i(t) > η(t)(Fig 4C and D). In other words, the larger the retinal slip was, the greater the input depolarising step current. These depolarising step current input stimuli, combined with the electrical coupling amongst IO cells regulated by inhibitory NO connections, generated the overall activity in the olivary system. Each individual CF spike conveyed well-timed information about the instantaneous error (See Fig 4D and E). The probabilistic spike sampling of the error ensured a proper representation of the entire error range across trials whilst maintaining CF activity between 1 and 10 Hz per fibre, which is consistent with electrophysiological data [22]. Even at this low frequency, it accurately sampled the error evolution [34, 35, 69-71].

#### Medial vestibular nuclei (MVN)

200 MVN cells were modelled as LIF neurons, divided into two groups of 100 cells each, corresponding to agonist and antagonist ocular muscles. Each MVN cell received inhibitory input from a PC and excitatory input from the CF, which simultaneously innervated the corresponding PC. Additionally, each MVN cell received excitatory projections from all MFs, maintaining the baseline activity of MVN cells. The spike activity of both the agonist and antagonist groups of MVN cells was translated into an analogue output signal (eye velocity) according to equations 5 and 6:

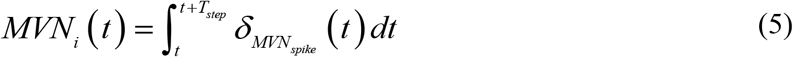

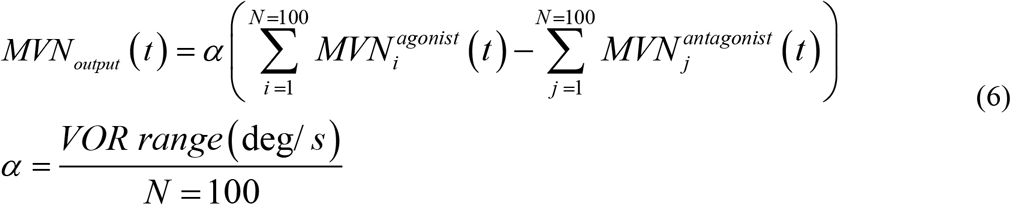

where α represents the kernel amplitude that normalises the contribution of each MVN cell spike to the cerebellar output correction. i and j are used to represent the MVN neuron tags, ranging from one to N = 100, which is the total number of MVNs in each sub-population (both agonist and antagonist sub-populations).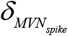 stands for the Dirac delta function that represents MVN spikes that have been triggered, while *T*_*step*_ (0.002 seconds) corresponds to the duration of the sliding windows over which the MVN spiking activity is calculated. This neural topology is summarised in Table1.

### D. Neuron Models

#### 1. The LIF model

The LIF model used for MFs and GCs was the same as the one used in [27]. However, the LIF model used for MVN was implemented based on [25] following equations 7-13. The neural dynamics of MVN were defined by the membrane potential and the presence of excitatory (AMPA and NMDA) and inhibitory (GABA) chemical synapses.

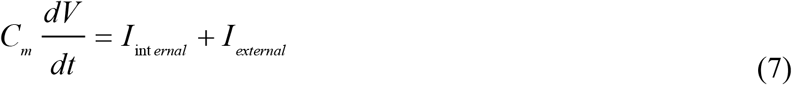

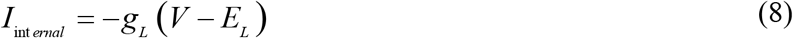

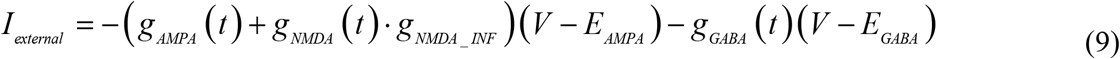

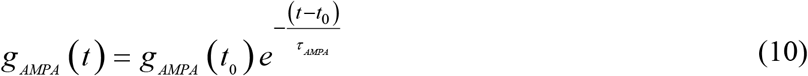

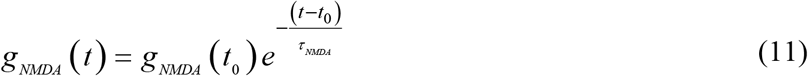

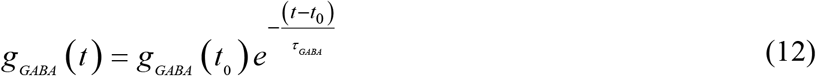

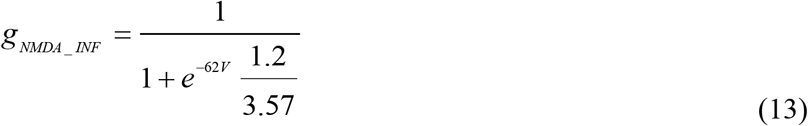

where C_m_ denotes the membrane capacitance, V the membrane potential, I_internal_ the internal currents and I_external_ the external currents. E_L_ is the resting potential and g_L_ the conductance responsible for the passive decay term towards the resting potential. Conductances g_AMPA_, g_NMDA_ and g_GABA_ integrate all the contributions received by each receptor type (AMPA, NMDA, GABA) through individual synapses. These conductances are defined as decaying exponential functions [47, 72, 73]. Finally, *g*_*NMDA_INF*_ stands for the NMDA activation channel.

#### *2*. The IO HH model

The IO model was a simplified and corrected version of the three-compartment HH model proposed by [19]. To enhance computational performance, a simplified set of somatic and dendritic currents was adopted, whilst still preserving the ability to generate spike bursts due to the sodium current inactivation within the axon hillock compartment [74]. Initially, the model was implemented in NEURON for validation in isolation, and subsequently transferred to EDLUT to accelerate the computation of the entire network.

**SOMA:** The total soma voltage was given by:

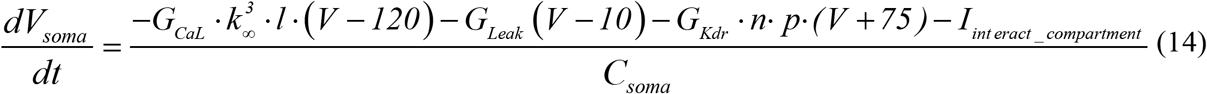

Where *C*_*soma*_ is the soma membrane capacitance and the dynamics of each gating variable follows:

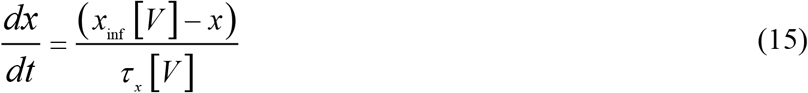

The equilibrium function *x*_*inf*_ [*V* ] and time constant for each current are depicted in table 2

**Table 2.**
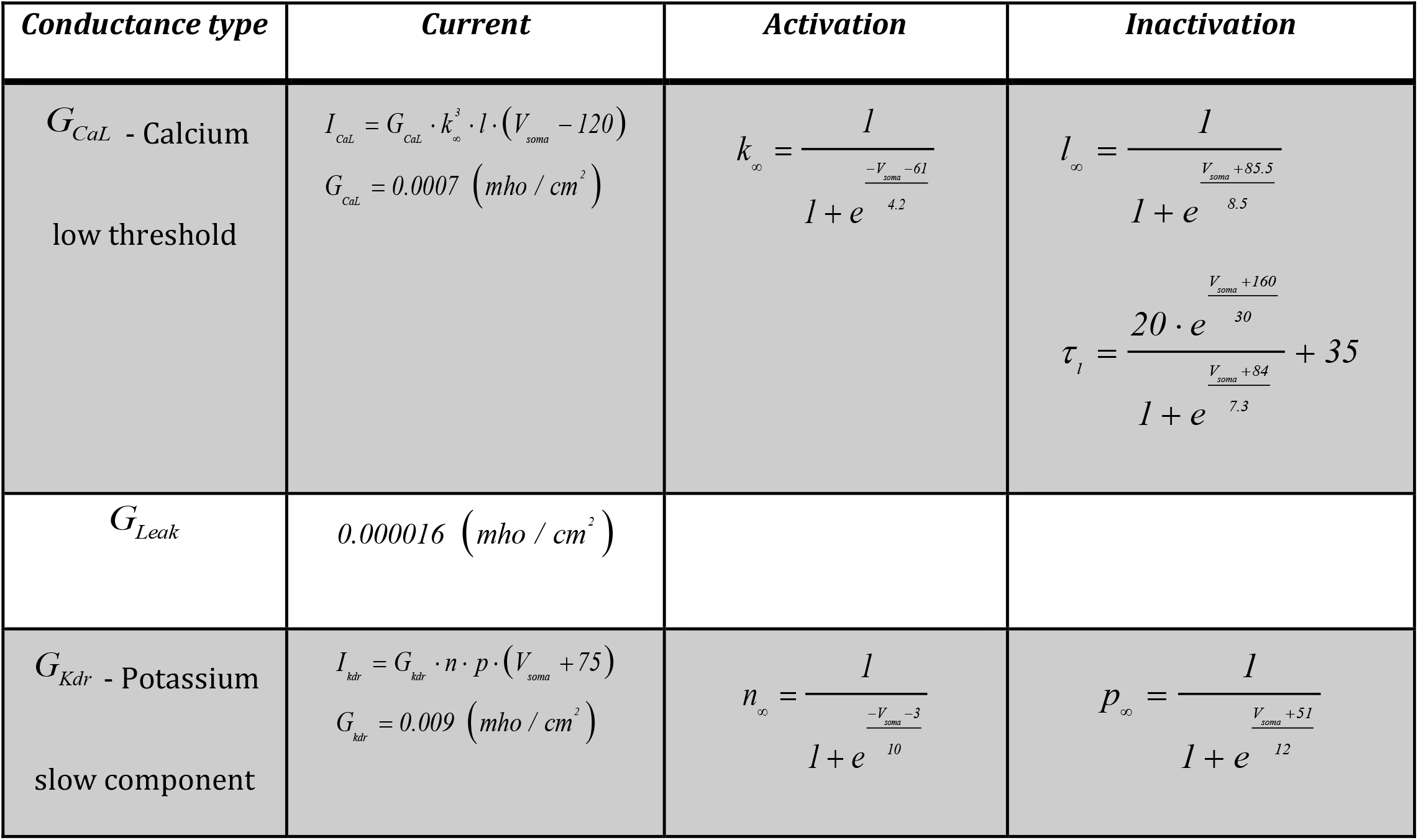
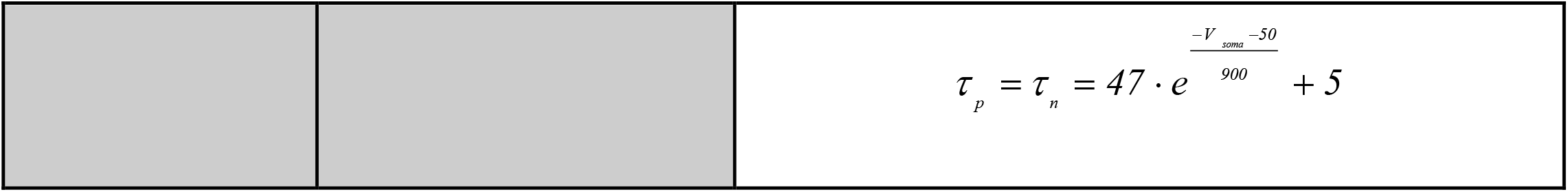
Somatic component. Ionic conductance kinetic parameters.

**AXON HILLOCK:** The total axon voltage was given by:

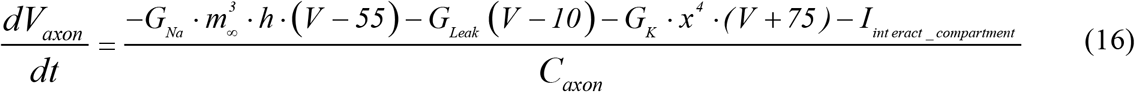

Where *C*_*axon*_ is the axon membrane capacitance and the dynamics of each gating variable follow Eq. (15). The equilibrium function *x*_*inf*_ [*V* ] and time constant for each current are depicted in table 3:

**Table 3.**
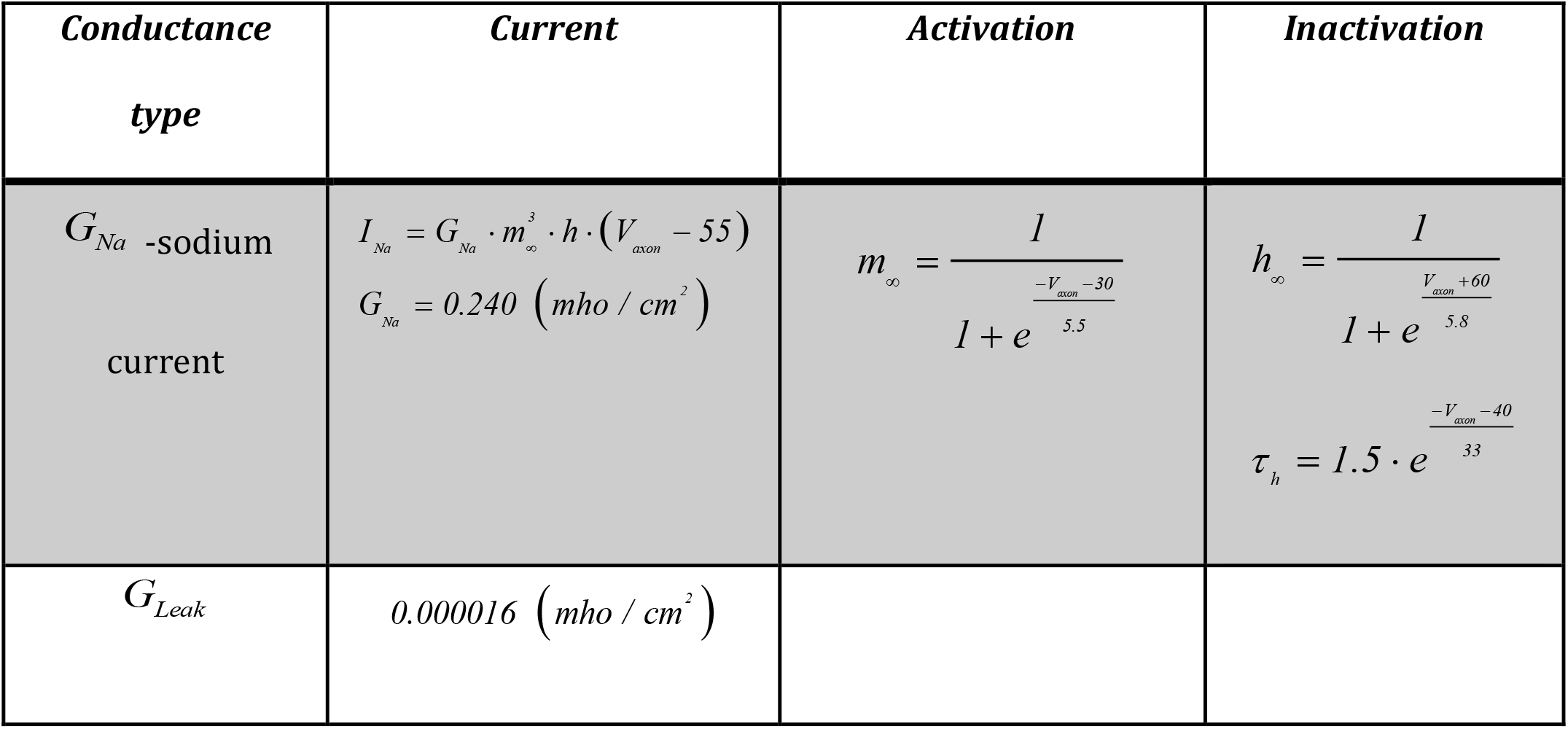
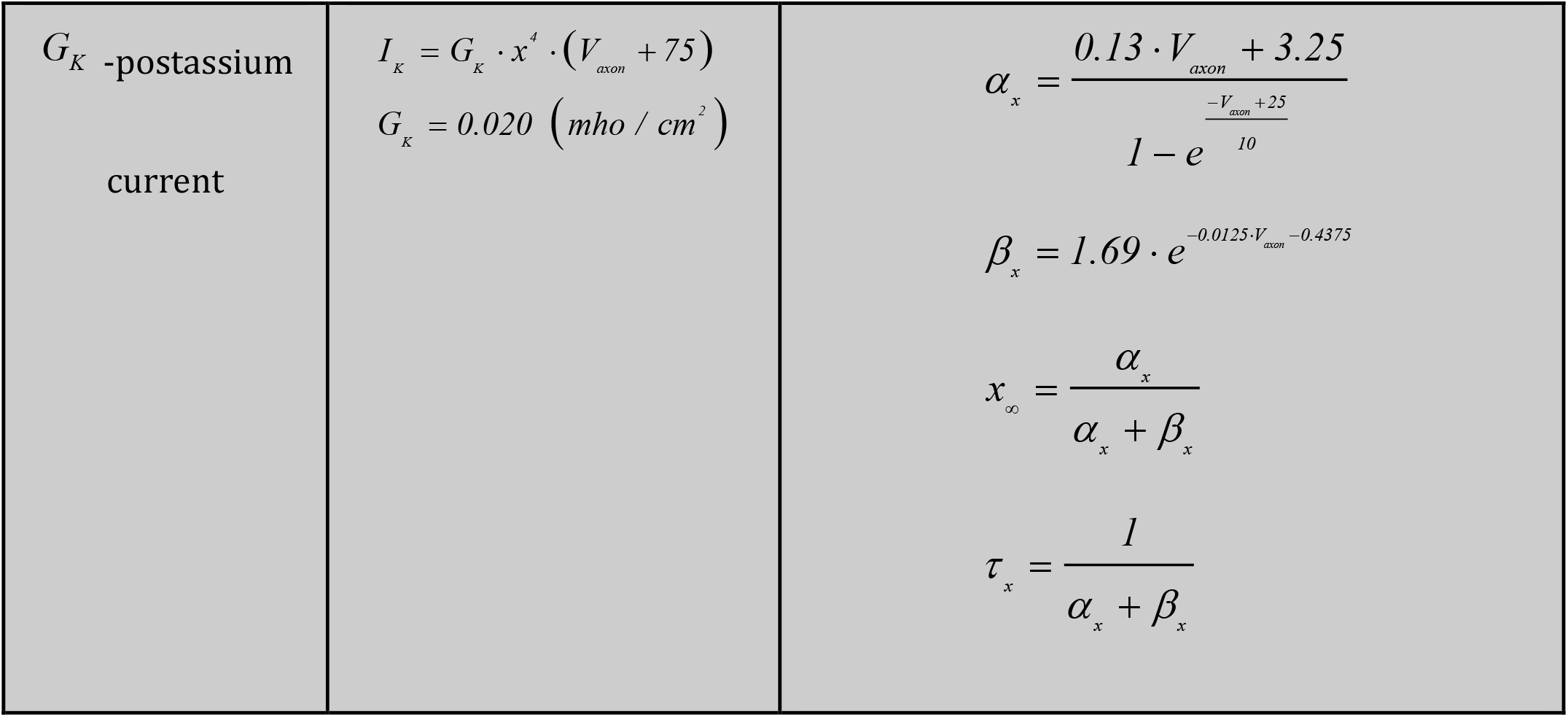
Axon component. Ionic conductance kinetic parameters.

| *Conductance*<br>*type* | *Current* | *Activation* | *Inactivation* |
| --- | --- | --- | --- |
| $G_{Na}$ -sodium<br>current | $I_{Na} = G_{Na} \cdot m_{\infty}^3 \cdot h \cdot (V_{axon} - 55)$ $G_{Na} = 0.240 \text{ (mho / cm}^2\text{)}$ | $m_{\infty} = \frac{1}{1 + e^{\frac{-V_{axon} - 30}{5.5}}}$ | $h_{\infty} = \frac{1}{1 + e^{\frac{V_{axon} + 60}{5.8}}}$ $\tau_h = 1.5 \cdot e^{\frac{-V_{axon} - 40}{33}}$ |
| $G_{Leak}$ | $0.000016 \text{ (mho / cm}^2\text{)}$ | | |

|  |  |  |
| --- | --- | --- |
| $G_K$ -postassium<br>current | $I_K = G_K \cdot x^4 \cdot (V_{axon} + 75)$ $G_K = 0.020 \text{ (mho / cm}^2\text{)}$ | $\alpha_x = \frac{0.13 \cdot V_{axon} + 3.25}{1 - e^{\frac{-V_{axon} + 25}{10}}}$ $\beta_x = 1.69 \cdot e^{-0.0125 V_{axon} - 0.4375}$ $x_\infty = \frac{\alpha_x}{\alpha_x + \beta_x}$ $\tau_x = \frac{1}{\alpha_x + \beta_x}$ |

**DENDRITE:** The total dendrite voltage was given by

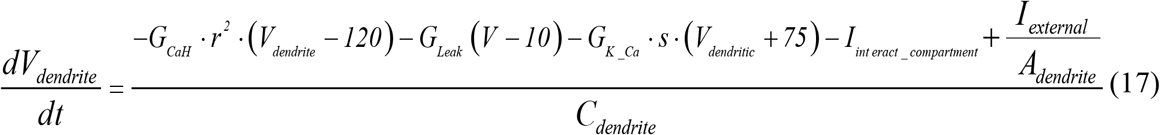

Where *C*_*dendrite*_ is the dendrite membrane capacitance, *A*_*dendrite*_ is the dendrite membrane area, and the external current *I*_*external*_ was given by:

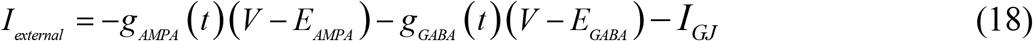

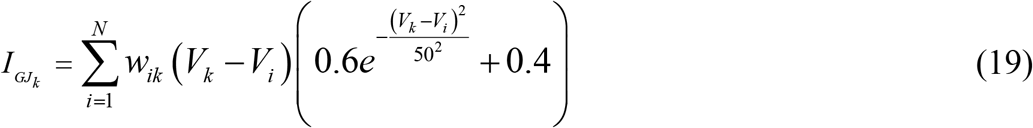

Where conductances gAMPA, and gGABA integrate all the contributions received by each receptor type (AMPA, GABA) through individual synapses Eq. (10, 12). Where 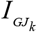 stands for the current injected to the *k*^*th*^ target neuron through the gap-junction (GJ) [20, 25, 28], *V*_*k*_ is the target neuron membrane potential, the i neuron membrane potential, *V*_*i*_ is the synaptic weight between the neuron i and the target neuron, and *N* is the total number of *GJ* current inputs. The dynamics of each gating variable follows Eq. (15). The equilibrium *x*_*0*_ [*V* ]?function and time constant for each current are depicted in table 4:

**Table 4.**
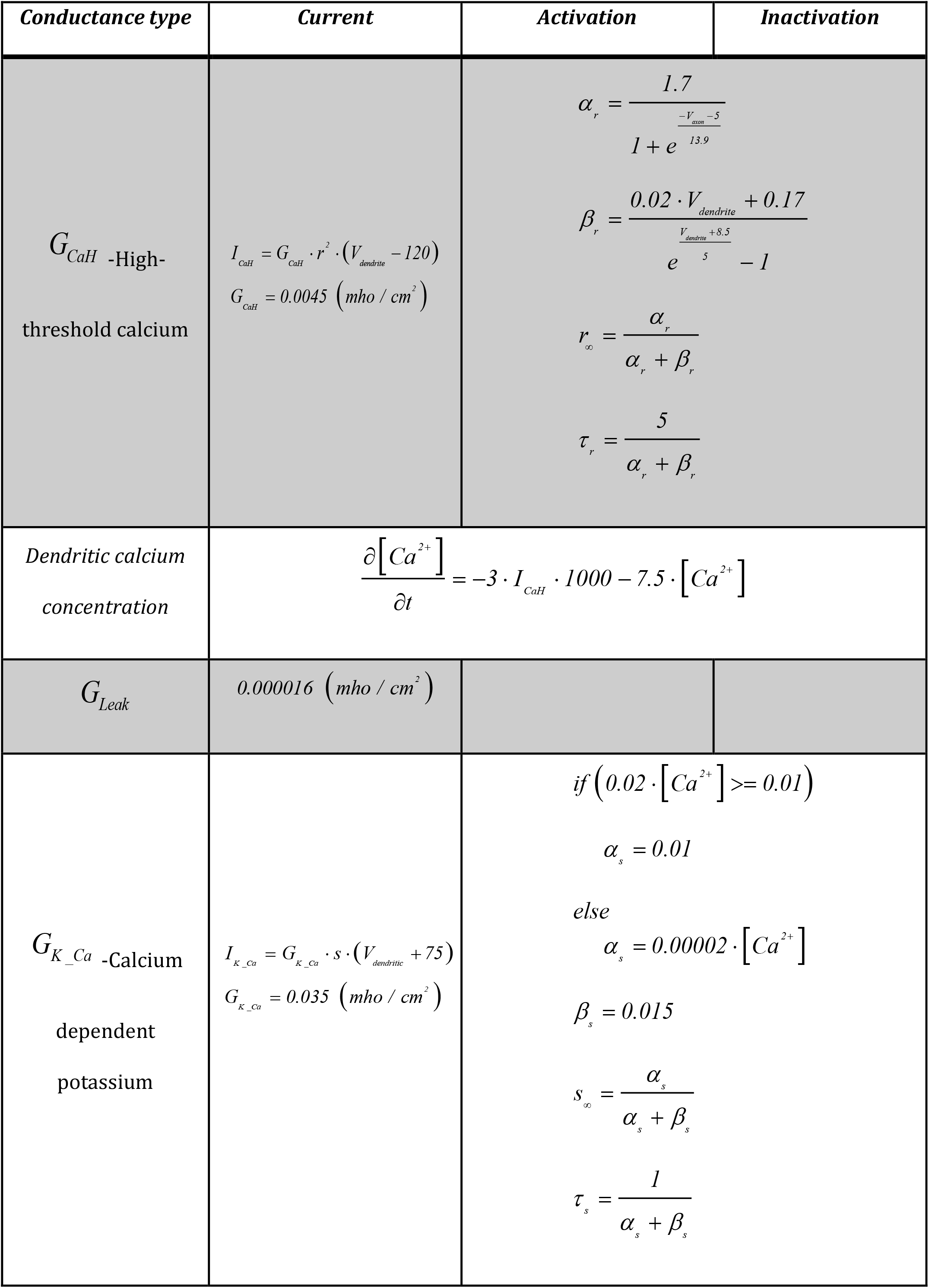
Dendritic component. Ionic conductance kinetic parameters.

**Table 5.** Geometrical parameters:

| **Soma Geometrical Parameters** |  |
| --- | --- |
| Cylinder length of the soma | **20.5790 $\mu m$** |
| Diameter of the soma | **20.5790 $\mu m$** |
| Membrane Capacitance | **1 $\mu F / cm^2$** |

| **Axon Geometrical Parameters** |  |
| --- | --- |
| Cylinder length of the axon | **53.1329 $\mu m$** |
| Diameter of the axon | **53.1329 $\mu m$** |
| Membrane Capacitance | **1 $\mu F/cm^2$** |
| **Dendrite Geometrical Parameters** |  |
| Cylinder length of the axon | **10.2895 $\mu m$** |
| Diameter of the axon | **10.2895 $\mu m$** |
| Membrane Capacitance | **1 $\mu F/cm^2$** |

The interaction between compartments was modelled passively taking the surface ratio between compartments into account.

Soma to dendrite current:

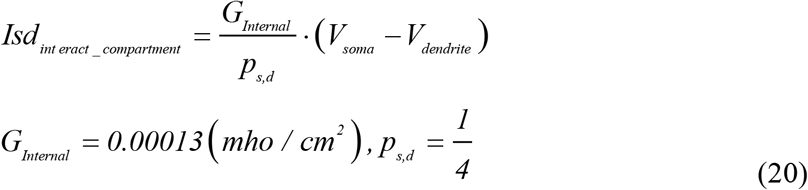

Axon to soma current:

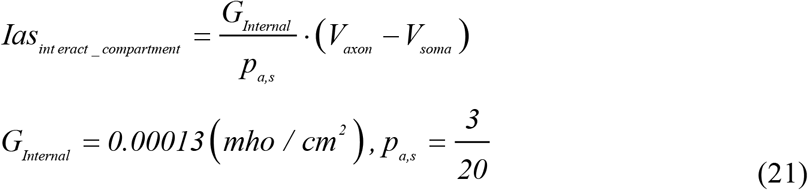

Soma to axon current:

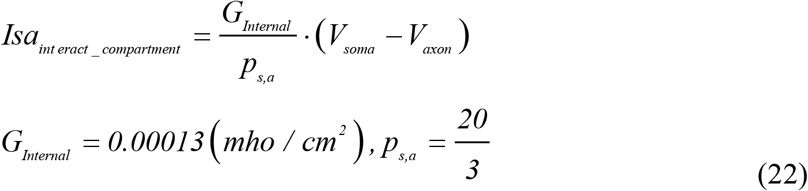

#### *3*. The PC HH model

The HH single-compartment model (PC) was based on [75, 76] and implemented in [27]. It consisted of a single compartment HH neuron with five ionic currents and excitatory (AMPA) and inhibitory (GABA) chemical synapses:

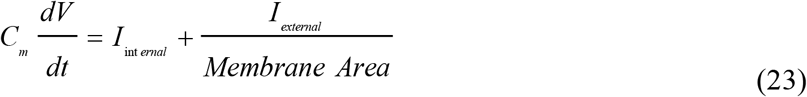

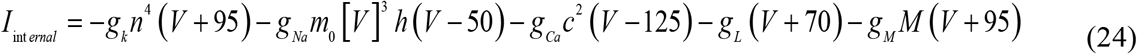

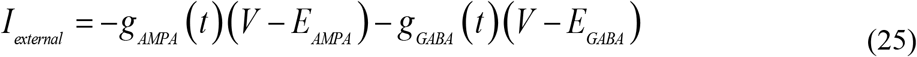

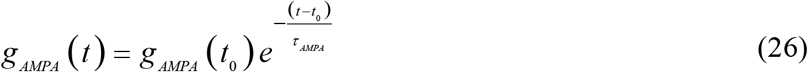

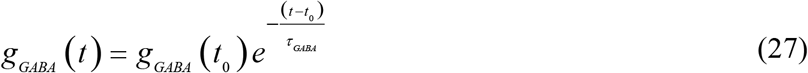

where *V* denotes the membrane potential, *I*_*internal*_ the internal currents and *I*_*external*_ the external currents. *C*_*m*_ is the membrane capacitance. Conductances *g*_*AMPA*_ and *g*_*GABA*_ integrate all the contributions received by each chemical receptor type (AMPA and GABA) through individual synapses as in [25, 27, 28]. These conductances are defined as decaying exponential functions. Finally, *g*_*K*_ is a delayed rectifier potassium current, *g*_*Na*_ a transient inactivating sodium current, *g*_*Ca*_ a high-threshold non-inactivating calcium current, *g*_*L*_ a leak current, and *g*_*M*_ a muscarinic receptor suppressed potassium current. The dynamics evolution of each gating variable (*n, h, c*, and *M*) can be computed using Eq 15. Where *x* indicates the variables *n, h, c*, and *M*. Gating variables are defined in [27] .

### E. Synaptic Plasticity

The overall input-output function of the cerebellar network model incorporated two STDP mechanisms at different sites, which balanced long-term potentiation (LTP) and long-term depression (LTD). For a more detailed review of the implemented synaptic mechanisms, refer to [25, 27, 28, 35].

#### 1. PF–PC synaptic plasticity

The LTD/LTP balance at PF–PC synapses is based on:

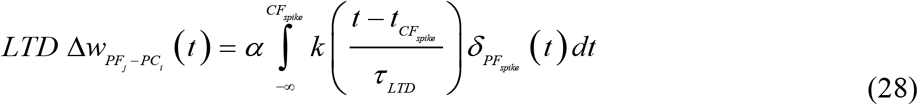

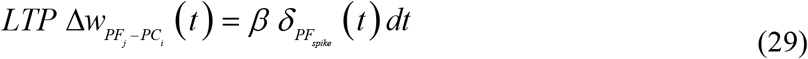

where Δ*W*_*PFj–PCi*_*(t)* denotes the weight change between the *j*^*th*^ PF and the target *i*^*th*^ PC; τ_LTD_ = 100 ms denotes the time constant that compensates for the sensorimotor delay; *δ*_*PF*_ is the Dirac delta function corresponding to an afferent spike from a PF; *α* = -0.0304 nS is the synaptic efficacy decrement; *β* = 0.0184 nS is the synaptic efficacy increment; and the kernel function *k(x)* [25, 27-29] is defined as:

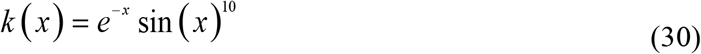

The STDP mechanism, as described in (Luque, et al., 2016), results in synaptic efficacy decrement (LTD) when a spike from the CF reaches the target PC neuron. The extent of this synaptic decrement is determined by the activity arriving via PFs, which is convolved with an integrative kernel defined in Eq. (30) and then scaled by the synaptic decrement factor *α*. This effect on the presynaptic spikes arriving through PFs is most pronounced within a 100 ms window preceding the arrival of the postsynaptic CF spike. This temporal window compensates for the sensorimotor pathway delay [70, 77-79] .On the other hand, the amount of LTP at PF-PC synapses remains fixed, with each spike arriving through a PF to the targeted PC resulting in an increase in synaptic efficacy equal to *β*. In the simulated loop, The sensory-motor pathway delay [80], with a duration of 100 milliseconds, was modelled using two circular temporal buffers, each lasting 50 milliseconds and having 2-millisecond taps. The first buffer was positioned between the cerebellar output and the r-VOR plan, whilst the second buffer was situated between the output of the r-VOR plant and the error signal used as the cerebellar instructive signal (retinal slips) [25].

#### 2. MF–MVN synaptic plasticity

The LTD/LTP dynamics at MF – MVN synapses are based on:

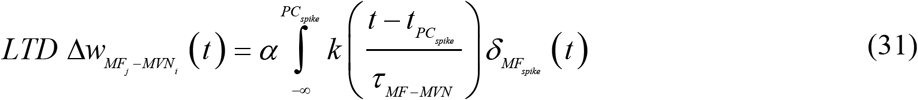

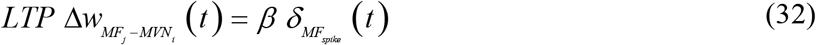

with Δ*W*_*MFj–MVNi(t)*_ denoting the weight change between the *j*^*th*^ MF and the target *i*^*th*^ MVN; *τ*_*MF-MVN*_ = 5 ms standing for the time width of the kernel; *δ*_*MF*_ representing the Dirac delta function that defines a MF spike; *α* = -0.002048 nS is the synaptic efficacy decrement; *β* = 0.000792 nS is the synaptic efficacy increment; and the integrative kernel function *k(x*) [25, 27-29, 35] defined as:

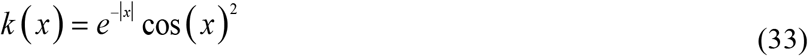

The STDP results in a synaptic efficacy decrease (LTD) when a spike from the PC reaches the targeted MVN neuron. The extent of this synaptic decrement is influenced by the activity arriving via MFs, which is convolved with the integrative kernel defined in Eq. (33) and then scaled by the synaptic decrement factor *α*. This LTD mechanism takes into account presynaptic/postsynaptic MF spikes that arrive before/after the postsynaptic/presynaptic PC spike within the time window defined by the kernel (*τ*_*MF-MVN*_). Conversely, the amount of LTP at MF-MVN synapses remains constant, with each spike arriving through an MF to the targeted MVN resulting in an increase in synaptic efficacy defined as β.

## Additional informatioN

### Author contributions

Conceptualisation: Niceto R. Luque, Eduardo Ros, Angelo Arleo

Data Curation: Niceto R. Luque, Francisco Naveros, Richard R. Carrillo

Formal Analysis: Niceto R. Luque, Angelo Arleo

Funding Acquisition: Niceto R. Luque

Investigation: Niceto R. Luque, Francisco Naveros, Richard R. Carrillo

Methodology: Niceto R. Luque, Angelo Arleo

Software: Niceto R. Luque, Francisco Naveros, Richard R. Carrillo

Supervision: Eduardo Ros, Niceto R. Luque, Angelo Arleo

Writing – Original Draft Preparation: Niceto R. Luque

Writing – Review & Editing: Niceto R. Luque, Eduardo Ros, Angelo Arleo

No competing interests declared by none of the authors.

